# A Myb-dominated gene regulatory network universally controls sexual cell fate transitions in diatoms

**DOI:** 10.64898/2026.03.25.714157

**Authors:** Gust Bilcke, Arthur Cleyman, Nadine Rijsdijk, Triana Forment, Thomas Eekhout, Darja Belišová, Peter Chaerle, Carolin Grones, Sien Audoor, Michiel Van Bel, Judith Porters, Nicolás Manosalva Pérez, Evelien Mylle, Daniël Van Damme, Bert De Rybel, Lieven De Veylder, Wim Vyverman, Klaas Vandepoele

**Author notes:** These authors contributed equally (Arthur Cleyman, Nadine Rijsdijk). These authors share senior authorship (Lieven De Veylder, Wim Vyverman, Klaas Vandepoele). Laboratory of Cell and Developmental Biology, Wageningen University, Wageningen, The Netherlands (Carolin Grones). Laboratory of Cell Cycles of Algae, Centre Algatech, Institute of Microbiology of the Czech Academy of Sciences, Trebon, Czech Republic (Sien Audoor).

## Abstract

Diatoms are the foundation of aquatic food webs and contribute about 40% of the total marine primary productivity. Yet, the regulation of their complex size-dependent life cycles remains obscure. Here, we leveraged single-cell transcriptomics and transgenic reporter lines to uncover the molecular mechanisms behind partner recognition, nuclear fusion, and the remarkable 15-fold size expansion of auxospores. Gene regulatory network inference revealed that the irreversible commitment to differentiate into gametes is controlled by Myb transcription factors, whose specific activity in the global ocean underscores their significance for ploidy transitions across diatom clades. These findings reinforce microalgae as powerful models to study cell fate transitions and provide a mechanistic framework for the life cycle dynamics that underpin the functioning of aquatic systems worldwide.

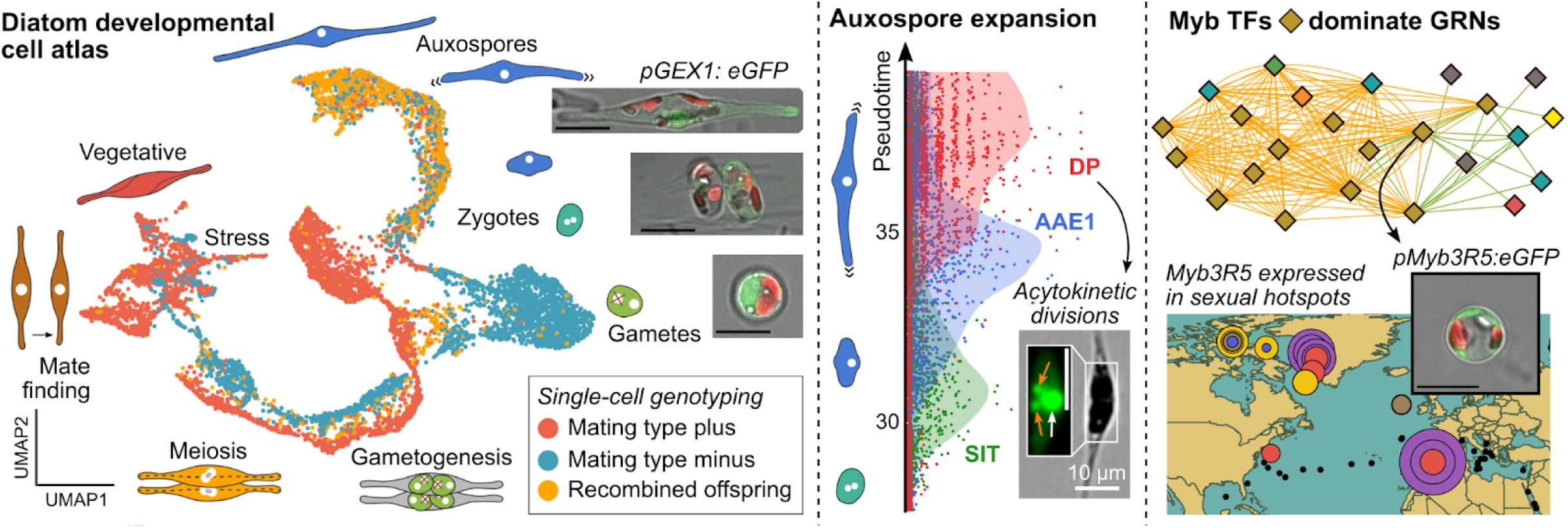

## Introduction

Sexual reproduction can be traced back to the last eukaryotic common ancestor 1.8-2.7 billion years ago ^1^ and is widely conserved across contemporary eukaryotic taxa ^2^. As gamete formation and fusion are crucial in transmitting genetic information to the next generation, multicellular organisms have evolved complex developmental systems that control gametogenesis, gamete recognition and embryogenesis. In contrast, unicellular eukaryotes (protists) have historically been regarded as predominantly reproducing via mitosis and only occasionally engaging in sex, controlled by simple environmental triggers ^3,4^. Yet, diatoms, the most species-rich eukaryotic algae that single-handedly produce 20% of the global oxygen supply ^5^, stand out by their complex and endogenously controlled life cycles. Unlike many other protist taxa, diatoms are diploid during most of their life cycle, with only a brief haploid stage during sexual reproduction^4^. During their evolution, they have transitioned from oogamy to isogamy, driving one of the largest species radiations in eukaryote history ^6–8^. Because vegetative cell division is constrained by their rigid silica cell wall, diatom populations experience a gradual cell size decrease that eventually results in cell death, but is counteracted by the size expansion of a unique but poorly understood auxospore ^9^. Despite the necessity of sexual reproduction for the survival of diatom populations, the transcriptional network that controls the irreversible sequence of cell fate transitions is completely unknown. Indeed, the inherent cellular heterogeneity in laboratory and field samples has prevented an accurate delineation of cell types and the assignment of genes to cell types using bulk approaches ^10–13^. This is further compounded by a lack of efficient transformation methods in sexual model diatoms, limiting the study of gene function. Single-cell transcriptomics offers a powerful solution to capture cell-to-cell variability, but while the technique is widely adopted to study multicellular tissues and development, it has so far only been applied to a handful of algae ^14–19^.

Here, we unlock microwell-based single-cell RNA-sequencing (scRNA-seq) for diatoms to investigate sexual cell fate transitions, adopting the pennate diatom *Cylindrotheca closterium* as a genomic model for life cycle research ^10,20^ (**Fig. 1a**). Cell fates were validated by combining scRNA-seq time series with transcriptional reporter lines, single-cell genotyping, imaging flow cytometry and pheromone treatments. Leveraging the single-cell resolution, we reconstruct the network of transcription factors (TFs) that governs cell type transitions and pinpoint the universal diatom regulators that drive ploidy transitions in the global ocean.

**Figure 1:**
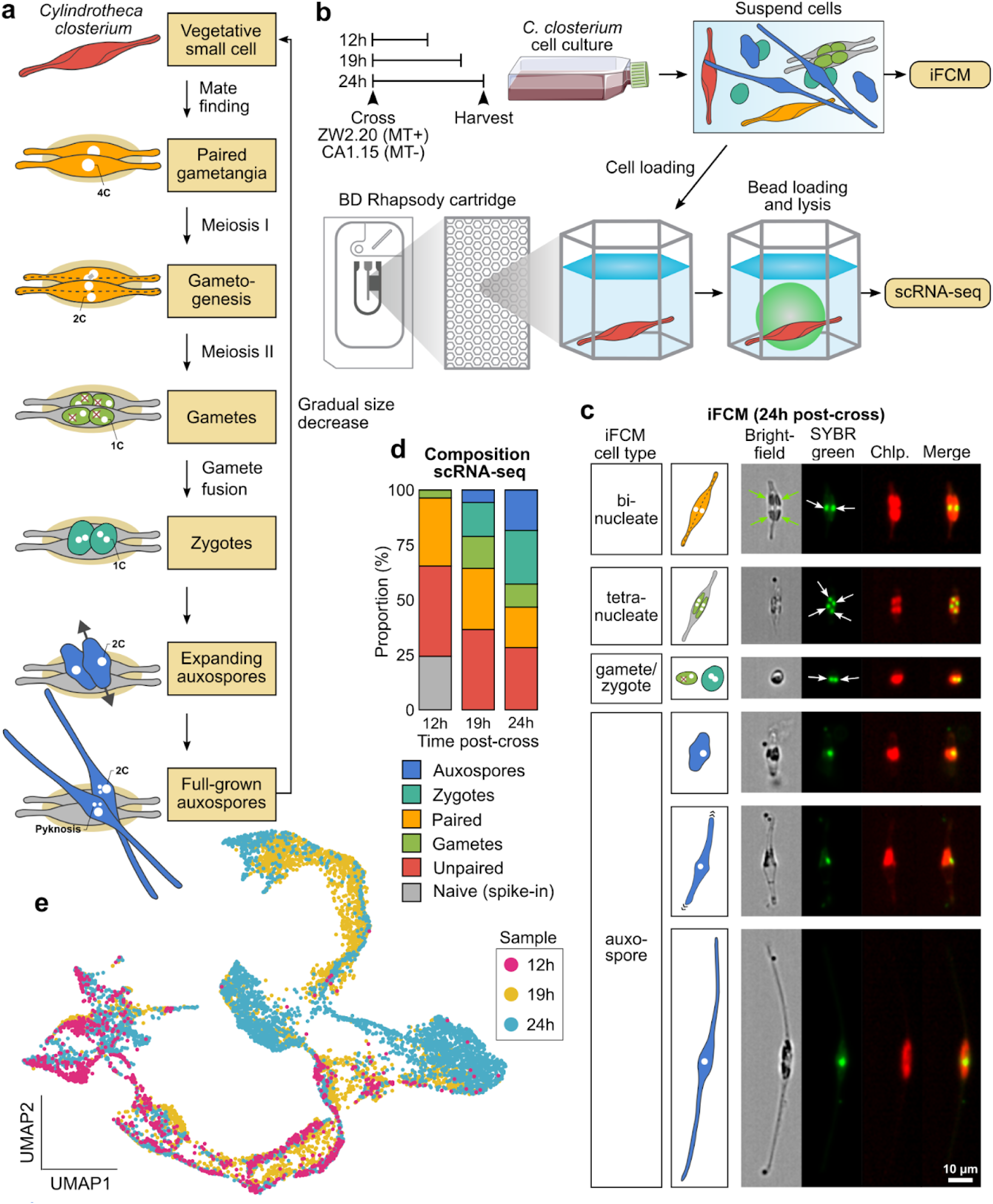
Experimental design of single-cell RNA sequencing (scRNA-seq) during life cycle transitions in the heterothallic diatom *Cylindrotheca closterium*. **(a)** Schematic representation of the sexual life cycle of *C. closterium*. The copy number of genetic material in the nucleus (white) in each cell type is indicated by “C”. Degradation of supernumerous haploid nuclei in gametes is indicated by red crosses. **(b)** Schematic overview of experimental workflow. CL = Cell Label. UMI = Unique Molecular Identifier. iFCM: imaging flow cytometry. **(c)** Microscopic pictures from the “24h” sample showing different cell types captured by an Amnis ImageStream X Mark II imaging flow cytometer. Images were obtained in brightfield, SYBR green and chloroplast autofluorescence (Chlp.) channels. The latter two were combined in the “Merge” channel. White arrows indicate individual nuclei while green arrows highlight the four chloroplasts of the binucleate gametangia stage. **(d)** Barplot showing the proportion of different cell types (colours) present at the three time points during the scRNA-seq experiment, as identified through light microscopy. **(e)** Uniform Manifold Approximation and Projection (UMAP) scatterplot of 8674 cells in the combined scRNA-seq dataset. Each cell is coloured by sample (time point).

## Results and Discussion

### Transient waves of expression drive a quick succession of sexual cell types

During sexual reproduction in the heterothallic diatom *C. closterium*, mating type plus (MT+, strain ZW2.20) is attracted towards mating type minus (MT-, strain CA1.15) to form a mating pair. Subsequently, both gametangia undergo meiosis and develop into two gametes each. After gamete fusion, the resulting zygotes expand until the auxospores each release a large-sized initial cell (**Fig. 1a**). In dark-synchronized laboratory cultures, this entire process takes about 24h (**Fig. S1**). To establish a timeline of sexual stages and their gene expression dynamics, we performed single-cell transcriptomics and imaging flow cytometry at three time points (12h, 19h and 24h) after crossing *C. closterium* (**Fig. 1b,c,S2, Text S1**). Each time point captured a unique mixture of cell types, enabling us to chart the entire process from start to finish (**Fig. 1d**). In total, scRNA-seq yielded 8674 high-quality single cells expressing 20,534 genes, representing 84% of all genes in the genome (**Fig. S3, Text S2**). Dimensionality reduction using UMAP (Uniform Manifold Approximation and Projection) revealed a developmental trajectory, the direction of which could be deduced from the contribution of each time point (**Fig. 1e,S4**) ^10^.

The seven morphologically distinguishable cell types resolved into 16 cell clusters, each defined by marker genes (**Fig. 2a,b,S5,S6, Text S3**). We next deployed a classifier based on known cell cycle genes to find that Cluster 1-4 cells (vegetative, mate finding) mostly resided in the G1-phase, while Cluster 5-9 (paired gametangia) underwent meiosis and cytokinesis (**Fig. 2c,S7,S8**). In terms of the number of marker genes, gametes retained a similar level of transcriptional activity as other cell types, as opposed to multicellular eukaryotes where gametes are typically transcriptionally dormant ^21–24^. Notably, gene expression changed gradually throughout sexual development, forming a continuum of intermediate states instead of discrete cell types. This prompted us to model expression in function of pseudotime by fitting a trajectory from start (vegetative cells) to finish (mature auxospores) (**Fig. 2d**). Anchoring pseudotime to the real developmental time revealed an almost perfectly linear relationship (adjusted R² = 0.99), indicating a relatively constant pace of transcriptional differentiation (**Fig. S9**). When examining trajectory-based markers, i.e. genes that were significantly upregulated at a specific pseudotime, we found that their expression only lasted for a short duration of time (**Fig. 2e**). Indeed, about half (48%) of all markers were active for less than four hours. This is reminiscent of the short-lived expression cascades that govern cell fate transitions during development of other eukaryotes, such as yeast spore germination ^25^ and animal embryogenesis ^26^.

**Figure 2:**
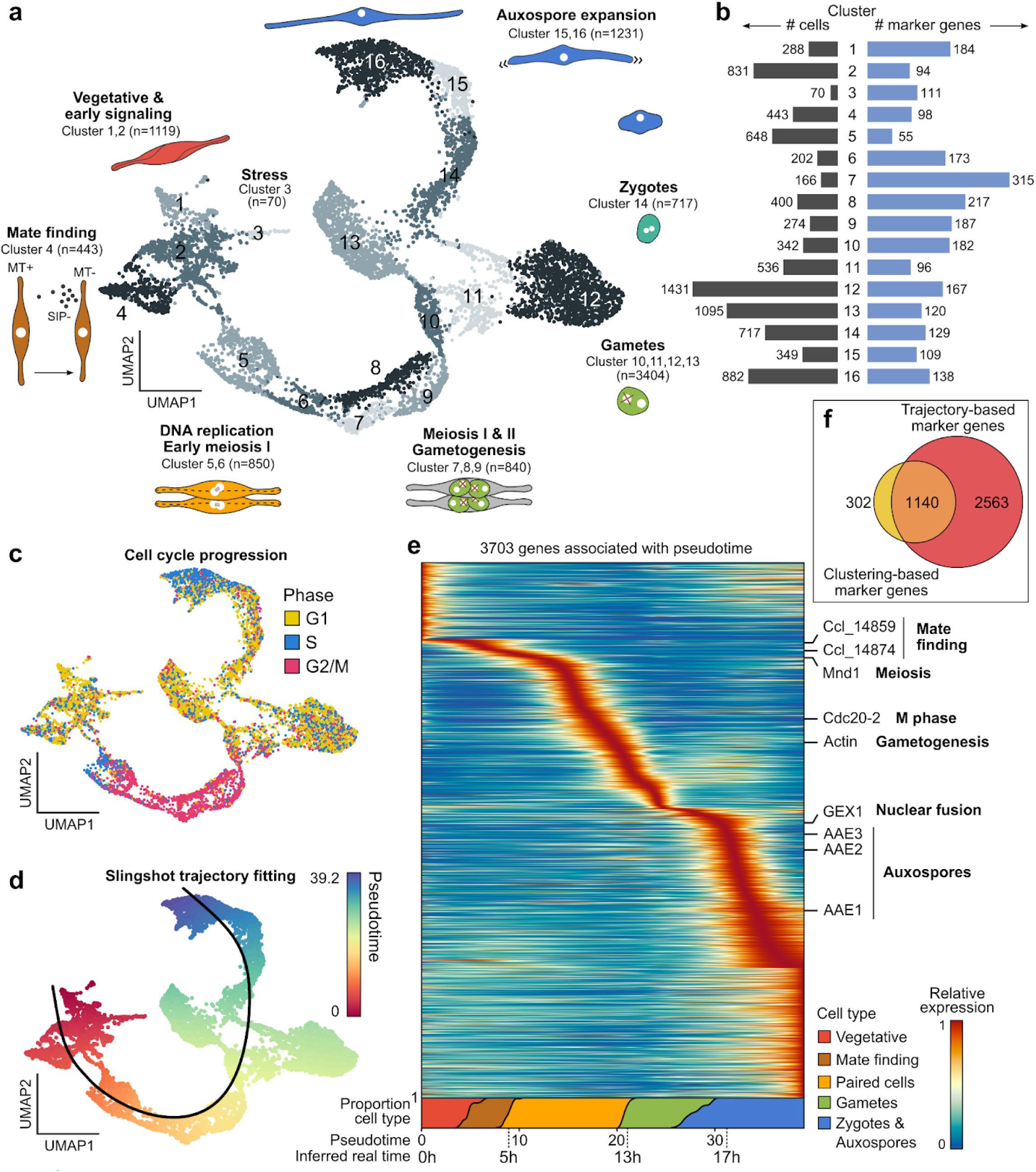
Identification of marker genes for diatom sexual cell types using clustering-based and trajectory-based approaches. **(a)** UMAP plot showing the subdivision of cells into 16 discrete clusters. Corresponding cell types are schematically shown at the outer edge of the plot, with “n” giving the total number of cells assigned to each cell type. **(b)** Bar plot showing the number of cells and cluster-based marker genes identified for each cluster. **(c)** UMAP plot showing the cell cycle phasing prediction for each cell (colors). **(d)** Trajectory curve superimposed on the scRNA-seq UMAP plot. Individual cells (dots) are coloured according to their projected pseudotime. **(e)** Heatmap of the relative expression of 3703 trajectory-based marker genes (rows) in function of pseudotime (x-axis). For each associated gene, the color scale shows gene expression as modeled by a generalized additive model (GAM) with eight knots, normalized to a maximum expression of one. Genes were ordered by their time of maximum expression. On the right, the position of a selection of marker genes with a known function is indicated. The inferred actual developmental time in hours since the start of a cross is shown below the pseudotime. AAE: Auxospore-Associated Expression, GEX1: Gamete Expressed Protein 1. **(f)** Euler diagram comparing the sets of clustering-based and trajectory-based marker genes.

Taken together, while going through seven cell fate transitions in a single day, diatom cells continuously differentiate at the transcriptional level, involving the transient expression of 4005 clustering- and trajectory-based marker genes (**Fig. 2f**). Only 43% of these markers could be detected using bulk RNA-seq ^10^, indicating that, apart from assigning genes to specific cell (sub)types, scRNA-seq enables identification of previously undetectable genes.

### Mating type-biased genes undergo a two-step activation

Taking advantage of allelic variation in mRNA reads to discriminate between the two mating types and their recombined offspring, we projected genotype identity onto the UMAP plot, revealing a clear separation between MT+ and MT- during pair formation and meiosis (Cluster 5-6), gametogenesis (Cluster 7-9) and gamete development (Cluster 10-13) (**Fig. 3a, Video S1**). Despite being transcriptionally differentiated, both mating types go through the same developmental stages, except for Cluster 4, an MT+ specific cluster for which we could not identify a corresponding MT- counterpart. To validate the identity of this cluster, we used bulk RNA sequencing of MT+ cells exposed to medium containing the MT- sex-inducing pheromone (SIP-), confirming that Cluster 4 consists of pheromone-conditioned cells (**Fig. 3b,c, S11**). Crucially, Cluster 4 cells expressed two myosin motor proteins implicated in diatom gliding motility (CaMyoB and CaMyoC) ^27^, pointing towards a transcriptional control of the chemotactic response towards SIP-.

**Figure 3:**
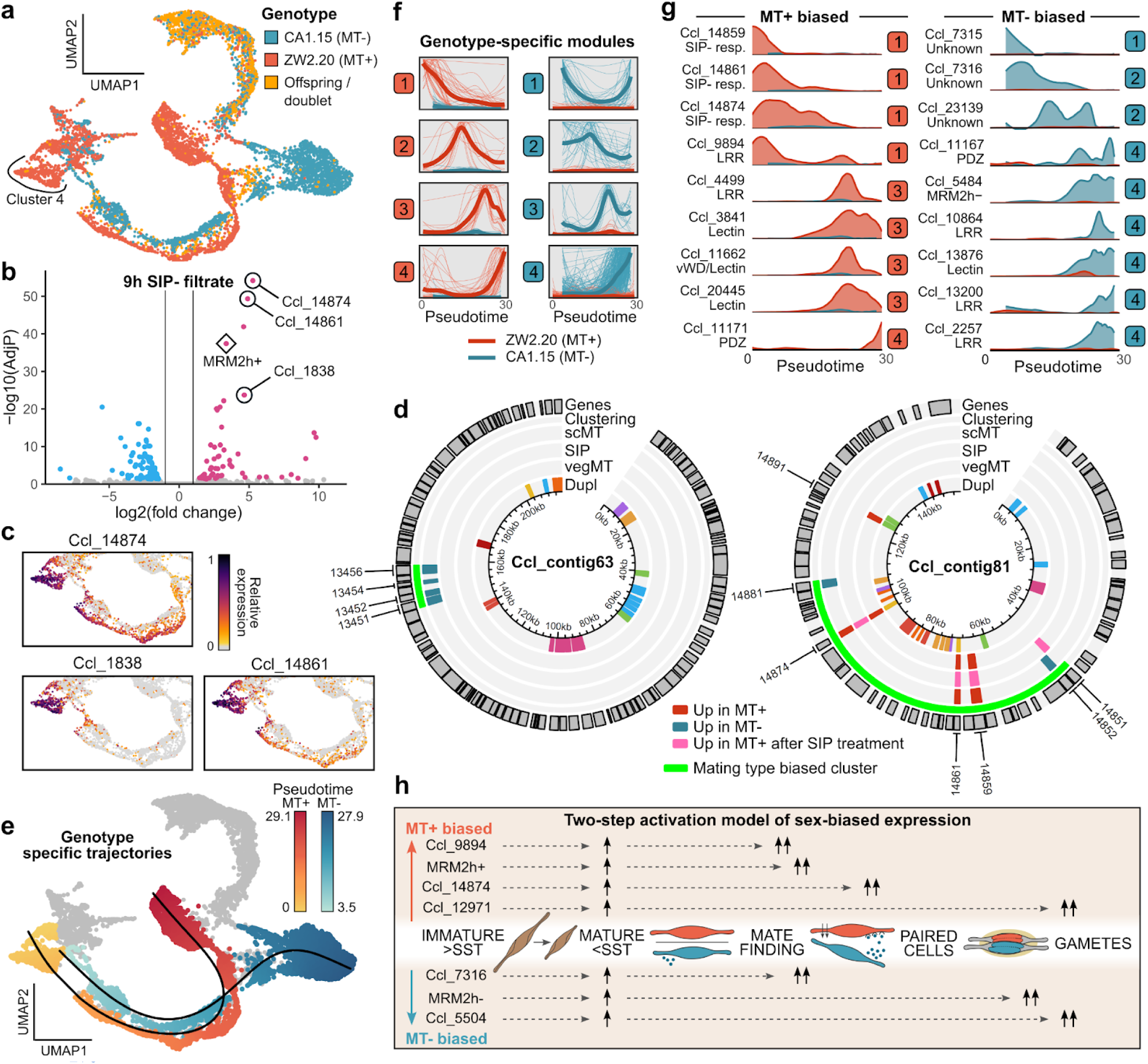
Differentiation between genotypes during sexual development. (**a**) UMAP plot showing the genotype of single cells. MT: mating type. (**b**) Volcano plot indicating the significance level of genes after 9h of sex-inducing pheromone minus (SIP-) treatment in function of their fold change. Significantly up- (pink) or downregulated (blue) genes are coloured accordingly. The top-3 marker genes for Cluster 4 in the single-cell RNA-seq data are highlighted with a circle. The known mating type-associated gene MRM2h+ (Mating-type Related Minus 2 plus homolog ^20^) is indicated with a diamond (**c**) Partial UMAP plots of the expression of the top-3 marker genes for Cluster 4. (**d**) Circos plots of two example contigs containing a mating type-biased gene cluster. Three types of differential expression are shown as concentric bands: between mating types in scRNA-seq data (scMT), in response to SIP and between mating types in vegetative cells of six different genotypes (vegMT) ^20^. Potential local duplicates belonging to the same gene family are indicated by colours in the inner circle (Dupl), while the identifiers of DE genes are shown on the outer edge. (**e**) UMAP plot showing two genotype-specific trajectories (lines) and the projected pseudotime of ZW2.20 (MT+) and CA1.15 (MT-) cells (coloured dots). Cells not used for trajectory fitting are shown in grey. (**f**) Relative expression (y-axis) over pseudotime (x-axis) of all mating type-specific genes divided into four modules for each mating type: [module 1] mate finding, [module 2] paired gametangia, [module 3] separating gametes and [module 4] mature gametes. Thick lines show the average expression over all genes while thinner lines represent individual genes. For each gene, the expression in MT+ (ZW2.20), and in MT- (CA1.15) is shown as separate lines in different colours. (**g**) Ribbon plots showing the Loess smoothed relative expression (y-axis) over pseudotime (x-axis) for a selection of genes. Expression of each genotype is separated by colours. (**h**) Proposed two-step model of mating type differentiation during stages of sexual reproduction. SST: sexual size threshold.

Following pheromone-guided attraction, MT+ and MT- gametangia establish a stable mating pair. This triggers differentiation into gametes, which will eventually fuse with the opposite mating type. To assess how this stage-wise process of partner recognition and fertilization functions, we performed a differential expression analysis between both mating types, revealing 482 highly specific mating type-biased genes. Despite the lack of sex chromosomes in diatoms ^28,29^, we did observe significant clustering of mating-type specific genes at six locations (**Fig. 3d, Fig. S12**). Modelling two parallel trajectories revealed a succession of four modules per mating type, corresponding to mate finding, paired gametangia, and separating and mature gametes, respectively (**Fig. 3e,f,S13**). Important for cell-cell recognition, each mating type expressed a sequence of leucine-rich repeat (LRR) proteins, including homologs of MRM2 (Mating-type Related Minus 2), a LRR protein implicated in partner recognition across pennate diatoms ^20,29^ (**Fig. 3g**). Meanwhile, several lectins were expressed in MT+ or MT- gametes, likely to facilitate cell-cell contact by binding carbohydrates on the compatible gamete, similar to lectin-mediated sperm-egg interactions in the related brown algae ^30–32^.

Interestingly, a significant portion of the genes that were pinpointed during cell-cell contact already showed a mating type-biased expression in vegetative Clusters 1 and 2 (Hypergeometric test, 6/8 modules p < 0.05, **Fig. S14**). Previous evidence shows that such mating type-specificity first originates upon sexual maturation, when vegetative cells decrease below a sexual size threshold ^20,29^ (**Fig. S15**). We therefore propose that sex determination in pennate diatoms follows a two-step activation, in which mating type-biased genes are primed when cells cross the sexual size threshold, and are further amplified during the stage of sexual reproduction when they are active (**Fig. 3h**). In conclusion, although gametangia and gametes of opposite mating types are morphologically indistinguishable, genotyping revealed a strong and dynamic transcriptional divergence between mating types underlying their behavioral differentiation and recognition.

### A molecular model of auxospore development captures nuclear fusion, cytoskeleton remodeling and acytokinetic divisions

We pinpointed the timing of gamete fusion to Cluster 14, because from this point onwards cells express genetic variants of both mating types (**Fig. 3a**). Each isogamete therefore seems to contribute transcripts to the zygote, which is immediately transcriptionally active, unlike plants and animals where the zygote is initially dominated by egg transcripts (“maternal control”) ^33,34^. Gamete fusion is followed by nuclear fusion, the timing of which was variable in *C. closterium*, occurring either in the zygote or early auxospores (**Fig. 4a**). Combining deep-homology searches, tertiary structure prediction and cross-species expression profiling, we uncovered the ancient nuclear fusogen Gamete Expressed Protein 1 (GEX1), which was considered to be absent from diatom genomes ^2,35^ (**Fig. S16, S17**). GEX1 reached its peak expression in young zygotes, but was already upregulated in gametes, a response previously reported in plants (**Fig. 4b**) ^36,37^. We confirmed this behavior by generating a transcriptional reporter line for GEX1, marking the first successful transformation of *C. closterium*. A fluorescent signal accumulated in zygotes and auxospores, but also in some of the gametes, supporting that an anticipatory expression of GEX1 precedes gamete fusion in diatoms (**Fig. 4c**).

**Figure 4:**
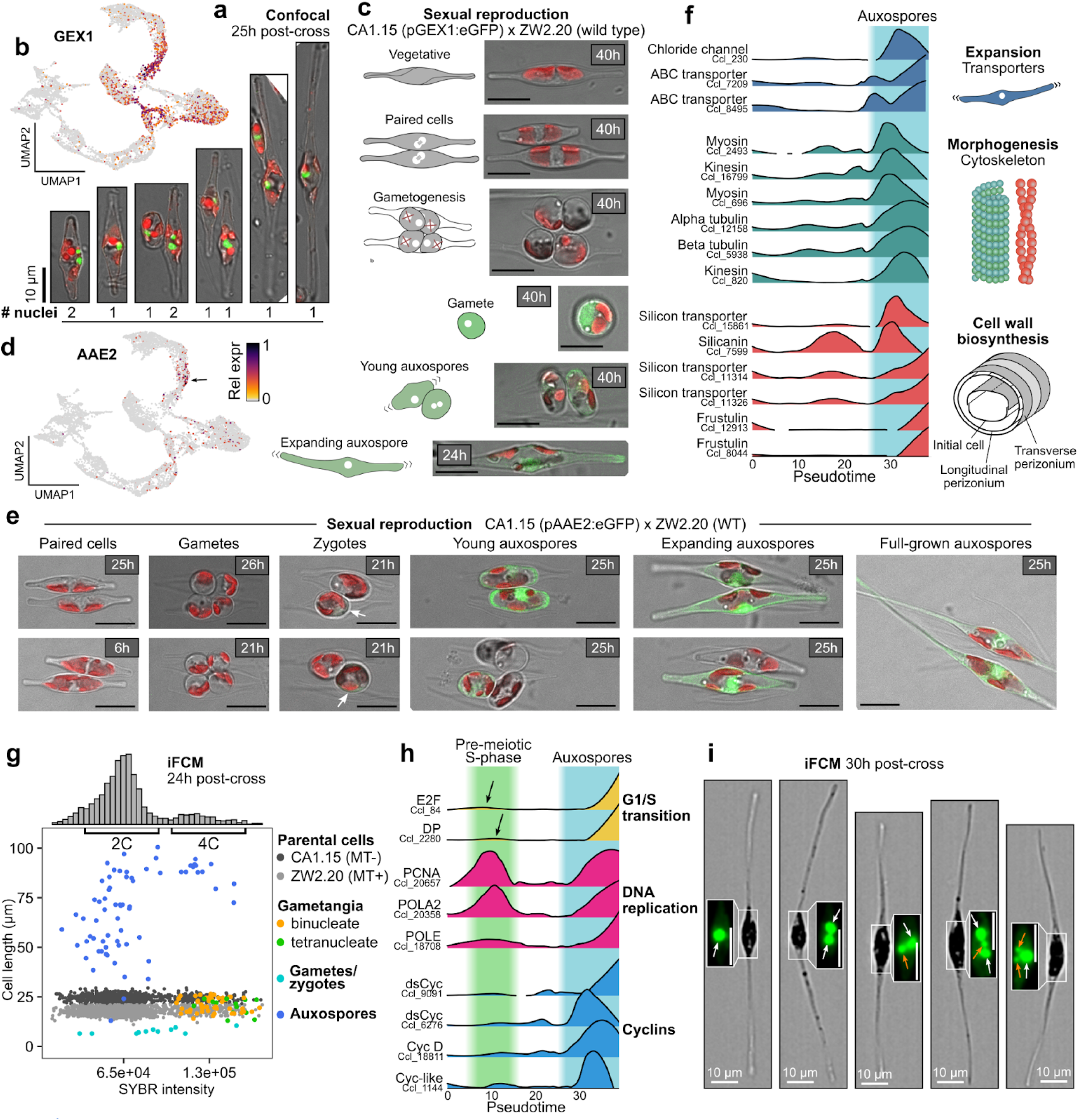
A molecular model of zygote and auxospore development. **(a)** Confocal microscopy images showing the variability in the timing of nuclear fusion during early auxospore expansion. Brightfield images are overlaid with nuclear SYBR Green staining (in green) and chlorophyll autofluorescence (in red). Scale bar: 10 µm. **(b,d)** UMAP plots of *C. closterium* cells coloured by the relative expression of *Gamete Expressed Protein 1 (GEX1)* and *Auxospore-Associated Expression 2 (AAE2)* respectively. **(c)** Confocal microscopy images of the *C. closterium* transcriptional reporter pGEX1:eGFP in different sexual cell stages. Red: autofluorescence, green: enhanced green fluorescent protein (eGFP). The time since crossing is shown in the upper right corner. Scale bars: 10 µm. **(e)** Confocal microscopy images of the *C. closterium* transcriptional reporter pAAE2:eGFP in different sexual cell stages. Red: autofluorescence, green: enhanced green fluorescent protein (eGFP). White arrows highlight the presence of a faint eGFP signal in zygotes. The time since crossing is shown in the upper right corner. Scale bars: 10 µm. **(f)** Ridgeline plot showing the density-smoothed expression of auxospore-expressed genes (y-axis) in function of pseudotime (x-axis). **(g)** Scatterplots showing SYBR intensity versus the total cell length, obtained by imaging flow cytometry (iFCM) from a subculture of the 24h scRNA-seq sample. Individual cells are coloured according to imaging-based cell type identification (nuclei, shape, ploidy, size). The histograms on top summarize the distribution of SYBR intensity of all cells. **(h)** Ridgeline plot showing the relative expression of cell cycle genes that are expressed during auxospore development, in function of pseudotime. Black arrows highlight a low level of expression during the pre-meiotic S-phase. DsCyc: diatom-specific cyclin **(i)** iFCM images of SYBR Green stained mature auxospores, 30h after crossing of CA1.15 and ZW2.20. White arrows: normal-sized nuclei. Orange arrows: pyknotic nuclei. Scale bars: 10 µm.

Following gamete fusion, zygotes with a diameter of ∼6.5 µm expanded into tubular auxospores with a length of up to 100 µm, representing a 15-fold increase (clusters 14-16, **Fig. S18**). To better understand the timing of auxospore development, we generated a reporter line for the AAE2 gene (Auxospore-Associated Expression 2)^10^, which was transiently expressed in Cluster 14 (**Fig. 4d**). A fluorescent signal became apparent in older zygotes and intensified after polarization, demonstrating that AAE2 marks the start of auxospore expansion, which enabled us to build a time-resolved molecular model of auxospore morphogenesis (**Fig. 4e**). Zygotes expressed several myosins, suggesting an actomyosin-mediated polarization mechanism like in brown algae ^38^. Next, a predicted vacuole-targeted ^39^ chloride-transporter marked early auxospore expansion, corroborating that the vacuole provides the osmotic force for expansion ^40^ (**Fig. 4f**). Auxospore expansion was accompanied by a strong and sustained expression of tubulins, while a cascade of cell wall-related genes supported the sequential formation of the zygote cell wall (incunabula), auxospore wall elements (perizonium) and initial cell wall (**Fig. 4f**) ^9,41^.

Even though auxospores never divide ^42^, we observed that (i) auxospores duplicated their genetic material when they reached a mature size of 80-100 µm and (ii) S-phase genes were reactivated during auxospore expansion (**Fig. 4g-h,S19**). By extending our time series to give auxospores more time to develop (30h post-cross), we could link these observations to acytokinetic divisions, a unique process in which the formation of each initial valve is preceded by a round of mitosis and nuclear degradation (**Fig. 4i**)^9^. Notably, several cyclins were specifically expressed during these acytokinetic divisions, making them early markers for the critical inflection point between sexual size restoration and the resumption of vegetative growth (**Fig. 4h**).

### A complex gene regulatory network regulates the diploid-to-haploid transition

The outspoken transcriptional dynamics that orchestrate cell fate transitions suggest that the diatom life cycle is under extensive regulatory control by TFs. Accordingly, 71 out of the 398 *C. closterium* TFs were markers for specific sexual cell stages. Even though heat shock factors are by far the most abundant TFs in the genome, Myb TFs, as well as Homeodomain, Cold Shock and Fork Head TF families were proportionally more involved in sexual reproduction (**Fig. 5a**). When only considering the TFs most significantly associated with pseudotime (AdjP < 1e-5), 32 TFs from 9 families remained, whose timing in development could be precisely pinpointed (**Fig. 5b**). The process of meiosis and gametogenesis was controlled by a quick succession of Myb TFs. Conversely, developing auxospores transcribed many different TF families, most notably C2H2, Heat Shock and Homeodomain TFs.

**Figure 5:**
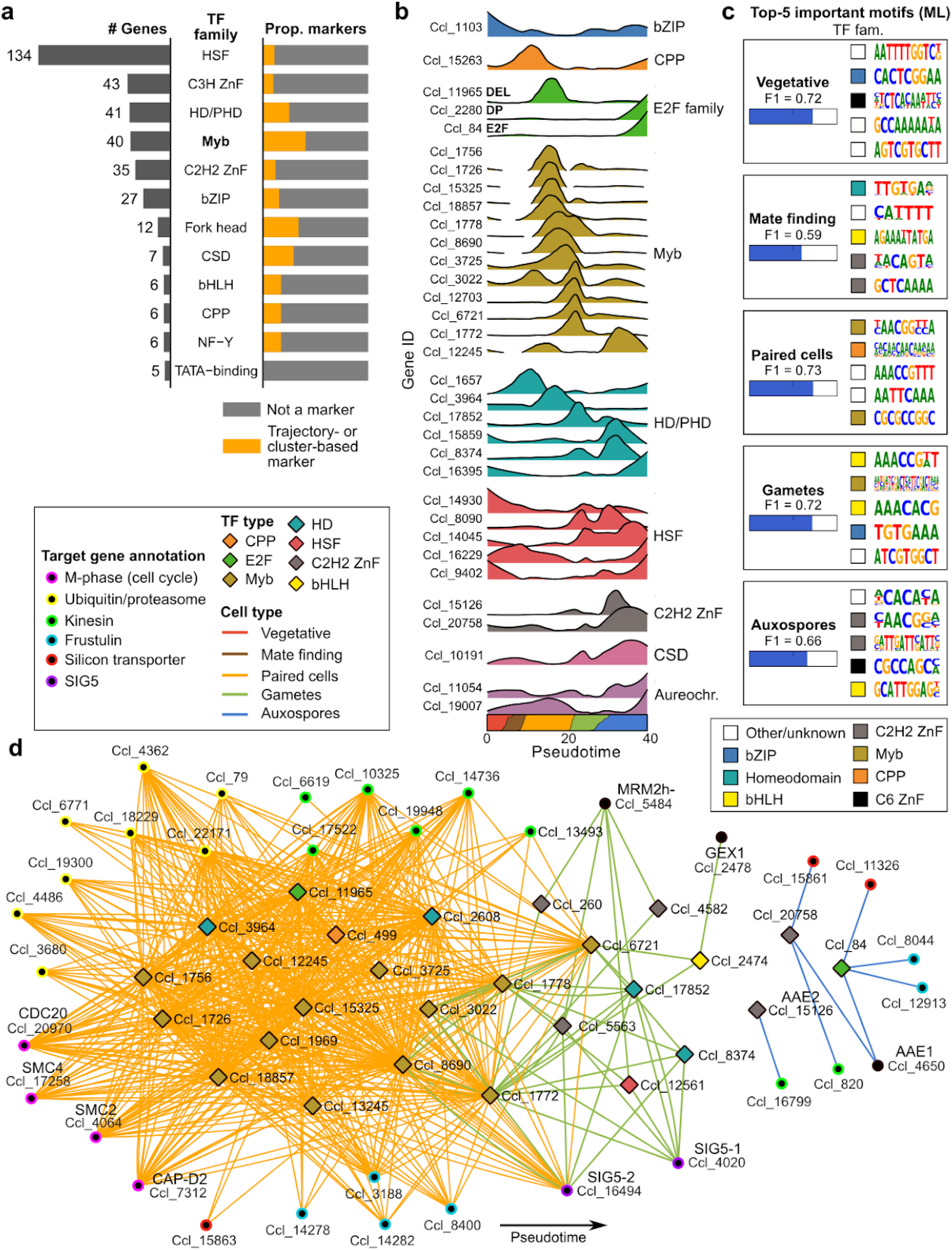
Gene regulation during sexual reproduction in diatoms. **(a)** Bar plots showing the number of genes and proportion of (clustering- and trajectory-based) marker genes among the most abundant transcription factor (TF) families in the *C. closterium* genome. **(b)** Ridgeline plots showing the density-smoothed relative expression of the top-32 associated TFs over pseudotime. TF families are indicated by colour. **(c)** Sequence logos of the top-5 most important motifs contributing to the prediction of the best performing machine learning (ML) model for each cell type. Coloured boxes (TF type hit) show the TF family that most likely binds each motif. **(d)** Gene regulatory network showing the predicted interactions (edges) between 27 transcription factors (diamonds) and a selection of relevant target genes (circles). Edges are coloured by the cell type where the regulon is active, diamonds are coloured by the transcription factor type and circles by the functional annotation of target genes. Transcription factors were approximately ordered by pseudotime along the x-axis. Abbreviations of known diatom sex genes: MRM: Mating-type Related Minus, SIG: Sex-Induced Gene, AAE: Auxospore-Associated Expression. Transcription factor family abbreviations: CPP: cysteine-rich polycomb-like protein, HD/PHD: homeodomain/plant homeodomain, HSF: heat-shock factor, ZnF: zinc finger, CSD: cold-shock domain. bHLH: basic helix–loop–helix.

Because the cis-regulatory code of diatoms is largely unknown, we trained machine learning models to identify sequence motifs whose occurrence in the promoter can predict gene expression within a sexual cell type (**Fig. 5c,S20,S21**). Several top ranked motifs resembled binding sites for well-known eukaryotic TFs and mirrored the expression of their corresponding TFs; with bZIP motifs driving transcription in vegetative cells, while CPP and Myb motifs predicted expression during meiosis and gametogenesis (**Fig. 5c,S22**). Next, we combined motif binding site information with single-cell co-expression to infer a gene regulatory network consisting of 7764 interactions and 946 genes, regulated by 27 unique TFs (**Fig. 5d**). The processes of meiosis and gametogenesis (Cluster 5-13) were strictly controlled, in particular by Myb TFs. Correspondingly, M-phase genes were enriched among the target genes of Myb TFs in Cluster 6 and 7 (hypergeometric test, AdjP < 0.05). Mybs also mutually activated each other in the network, probably explaining the quick succession of Myb TFs as an expression cascade (**Fig. S23**). Later in development, C2H2 and homeodomain TFs were predicted to drive the expression of putative gamete recognition genes such as *MRM2* and auxospore cell wall genes (**Fig. 5d**).

### Myb transcription factors govern diatom life cycle transitions across the globe

As our gene regulatory network pinpointed Myb TFs as central regulators of the diploid-to-haploid transition, we further investigated the Myb family in diatoms. Only a handful of Myb types were specific for sexual reproduction in *C. closterium*, in particular Myb3R5 and Myb2R2 (**Fig. 6a**) ^43^, the former being especially interesting because 3R Myb TFs are well-documented in cell cycle control of plants and animals ^44,45^. The Myb3R5 gene underwent repeated duplication in *C. closterium*, followed by expression divergence during meiosis and gametogenesis. Reporter lines for two “late” *C. closterium* Myb3R5 homologs never showed any eGFP fluorescence in vegetative growing cells, but a signal accumulated in gametes after crossing, confirming their role during the formation and maturation of haploid gametes from diploid gametangia (**Fig. 6b,c**). We assessed the functional conservation of sexual Myb TFs using bulk transcriptomics from two additional pennates (*Pseudo-nitzschia multistriata* and *Seminavis robusta*) and one centric diatom (*Skeletonema marinoi*). Across all species, Myb3R5 orthologs were on average 60-fold upregulated during gamete formation compared to vegetative growth (geometric mean). Pennate Myb2R2 homologs were also highly gametogenesis-specific, but their phylogenetic relationship with the centric Myb2R2 clade ^43^ could not be confirmed (**Fig. S24**).

**Figure 6:**
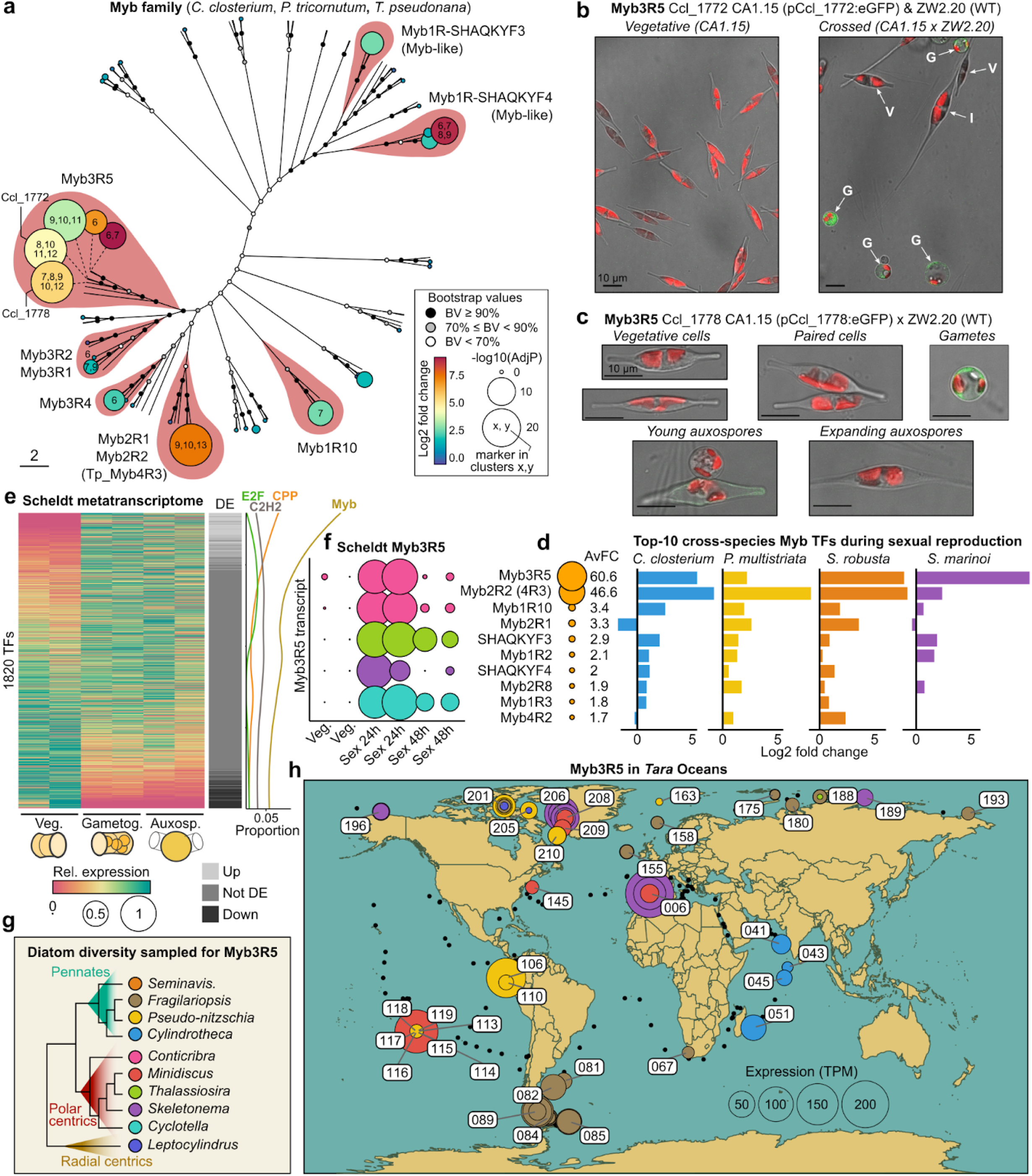
Myb transcription factors are central life cycle regulators in diatoms. **(a)** Equal-angle bootstrap-consensus phylogenetic tree of the Myb transcription factor (TF) family in three diatom species: *Cylindrotheca closterium*, *Phaeodactylum tricornutum* and *Thalassiosira pseudonana* (*Cyclotella nana)*. For each *C. closterium* gene, differential expression results during gametogenesis from bulk RNA-seq ^10^ are plotted as circles that scale with the -log10 false discovery rate-adjusted p-value (AdjP) and are coloured according to the log2 fold change. The clusters for which each Myb TF is a marker gene in the single-cell atlas are indicated with numbers inside the circles **(b)** Confocal images of the Myb3R5 reporter line Ccl_1772:eGFP in *C. closterium.* In the left panel, an uncrossed transgenic culture of CA1.15 is shown (negative control), while the right panel shows a crossed culture containing the following cell stages: V (vegetative small cell), G (gamete), I (initial cell). Red: autofluorescence, green: enhanced green fluorescent protein (eGFP). Scale bars: 10 µm. **(c)** Confocal images of the Myb3R5 reporter line Ccl_1778:eGFP in *C. closterium*. Scale bars: 10 µm. **(d)** Differential expression of Mybs during gametogenesis and zygote formation relative to vegetative growth, in four species: *C. closterium* ^10^, *Pseudo-nitzschia multistriata* ^13^*, Seminavis robusta* ^48^ and *Skeletonema marinoi* ^35^. Myb types were ordered by the average geometric mean of fold changes across species, as shown by circle radii. Bar plots show the log2 fold change during sexual reproduction for each species, averaged over paralogs where needed. **(e)** Heatmap showing the relative expression level and differential expression (DE) calls of 1820 Thalassiosirales TFs in metatranscriptome libraries where sexual reproduction was artificially induced using a salt-treatment ^46^. TFs are sorted from most upregulated (top) to most downregulated (bottom) during reproduction. Line plots visualize the Loess smoothed proportion of different TF families across the heatmap. The dominant cell type in each sample is drawn beneath the heatmap: vegetative (veg., control), gametogenesis (gametog., 24h) and auxospores (auxosp., 48h). **(f)** Bubble plot showing the expression of Myb3R5 homologs during the Scheldt microcosm experiment. The radius of circles varies with expression (counts per million, scaled to a maximum of 1 per transcript) and they are coloured by the genus that encodes the transcript (legend: see panel g). **(g)** Simplified cladogram showing the phylogenetic position of the diatom genera and species for which we investigated Myb3R5 expression in panel d,f and h. **(h)** World map highlighting the expression of Myb3R5 homologs in samples from the *Tara* Oceans expedition. Each circle represents an expressed Myb3R5 homolog in a single *Tara* sample, with the radius proportional to its abundance-normalized expression in that sample, as transcripts per million (TPM). Circles are coloured by the genus that is expressing a Myb3R5 homolog, following the colour scale in panel g. Myb3R5 transcripts expressed in the same station but from multiple genera / in independent MAGs / at different depths or size fractions were plotted superimposed. *Tara* stations where Myb3R5 was expressed are labelled with a station number. Black dots represent *Tara* stations where no Myb3R5 expression was detected.

To determine whether Mybs are also active as a life cycle regulator in natural ecosystems, we queried metatranscriptomes of an estuarine community where we artificially induced sexual reproduction in four genera of centric diatoms (*Cyclotella, Conticribra, Thalassiosira* and *Skeletonema*) ^46^. Among 1820 different TFs expressed by the Thalassiosirales, Mybs were about 3-fold more common among the top-100 upregulated TFs compared to non-responsive TFs (**Fig. 6e**). Each of the four sexualized genera expressed Myb3R5 with high specificity during gametogenesis (24h), demonstrating its potential to mark the diploid-to-haploid transition in the field (**Fig. 6f**). This prompted us to investigate the expression of Myb3R5 homologs encoded on metagenome-assembled genomes (MAGs) from the *Tara* Oceans expedition (**Fig. S25**). In total 17 MAGs from 7 diatom genera expressed Myb3R5, with the notable exception of the widespread genus *Chaetoceros*. Diatom Myb3R5 expression was observed in 36 *Tara* stations worldwide, and in particular at high latitudes where diatoms are most abundant ^47^ (**Fig. 6g,h**). Areas with predicted sexual activity ^46^ generally also showed Myb3R5 expression, including *Skeletonema* and *Minidiscus* near Gibraltar (station 006), *Pseudo-nitzschia* and *Minidiscus* in the South Pacific Ocean, and multi-species sexual hotspots near Greenland (stations 201, 205, 206). In line with this interpretation, the expression level of Myb3R5 was significantly correlated with sex signal strength in *Tara* Oceans samples (all t-tests *p* < 0.01, **Fig. S26**), supporting its function as a master regulator for life cycle transitions in the global ocean.

### Conclusions and perspectives

The incredible diversity of specialized and often ephemeral cell types found within a single water droplet has hampered our understanding of protist life cycles, which remain one of the most poorly appreciated aspects of biological oceanography ^49^. By introducing single-cell transcriptomics to track expression dynamics during life cycle transitions, we discovered that diatom genomes are tailored for life cycle regulation. Indeed, despite the lack of specialized reproductive tissues or embryogenesis programs, the number of cell clusters and marker genes expressed during sexual reproduction parallels that of multicellular animals ^50,51^ and plants ^52,53^, as well as that of the malaria parasite during its complex life history ^54–56^. As CRISPR-Cas has still not been achieved in any life cycle model diatom, we experimentally validated cell types using transgenic reporter lines, imaging, and single-cell genotyping. In doing so, we shed light on the molecular regulation of key life history features underlying the ecological success of diatoms ^6^, such as heterothallic partner recognition, the fusion of isogamous gametes and the remarkable process of size restoration.

Leveraging single-cell transcriptomics to construct gene regulatory networks is a powerful new approach to study the hundreds of TFs encoded in diatom genomes, of which only a tiny fraction has been functionally characterized. In diatoms, the diploid-to-haploid transition (meiosis, gametogenesis) is under tight regulatory control by Mybs, as well as E2F and CPP TFs, highlighting its importance as a critical decision point to balance the costs and advantages of sexual reproduction versus vegetative growth ^2,57^. In particular Myb3R5, whose role in diatom ploidy transitions seems to date back at least 160 Mya ^6^, showed widespread activity in the global ocean. This suggests that genome reduction and recombination is a frequent event in diatom populations, fostering the genetic diversity that is one of the root causes of the diatoms’ extraordinary species radiation, currently comprising ∼100,000 species ^58^, dwarfing all other taxa of phytoplankton combined ^59^.

Together with a simultaneous advance in single-nucleus RNA sequencing for the chain-forming diatom *P. multistriata* (Ruggiero et al., in prep.), the establishment of microwell-based scRNA-seq for protists is a key step towards capturing the functional diversity within natural phytoplankton and microphytobenthos assemblages. Therefore, the release of our integrative single-cell atlas as an interactive resource (www.single-cell.be/plants), will provide the community with access to this groundbreaking technology that has major repercussions for our understanding of ocean structuring, biochemistry and ecology ^49^.

## Material and Methods

### Induction of sexual reproduction in *Cylindrotheca closterium*

Two sexually compatible strains of *C. closterium* were obtained from the BCCM/DCG culture collection (https://bccm.belspo.be/about-us/bccm-dcg): CA1.15 as the MT- strain (DCG 0923) and ZW2.20 as its MT+ partner (DCG 0922). Cultures were grown in natural seawater supplemented with Guillard’s F/2 Marine Water enrichment solution (Sigma-Aldrich) and in the presence of antibiotics (0.7 g/L ampicillin, 0.5 g/L penicillin, 0.2 g/L streptomycin and 0.1 g/L gentamicin) to avoid bacterial growth. At the start of the experiment, cultures from both mating types were inoculated at a density of 2000 cells/mL and were grown in a 12:12 day:night rhythm for three days, after which medium was refreshed. On the fourth day, cells were suspended by gentle scraping and 15 mL of medium was transferred to small culture flasks (25 cm², VWR®), which were put in complete darkness for two days to synchronize the entry into sexual reproduction ^10^. Still in the dark, cells were resuspended by scraping and cultures of the compatible mating types were pooled at the same density, as determined using Burker Chamber counts under a Axiovert 40C inverted microscope. We then transferred 15 mL of suspended crossed cells to new culture flasks, which were moved into continuous light and a temperature of 21°C. Crossed cultures were allowed to develop for 12h, 19h or 24h before single-cell RNA sequencing (scRNA-seq). Just before loading, a spike-in of non-sexualized, naive cells of the ZW2.20 strain was added to the 12h sample, making up 24.4% of the final population. Just before loading the BD Rhapsody cartridge for scRNA-seq, we quantified the sexual cell stages present in each sample using an Axiovert inverted microscope at 20x magnification. Specifically, cells were scored as one of seven cell type categories using the cell counter plugin in FIJI (ImageJ): unpaired cells, paired cells, gametes, zygotes, young expanding auxospores, mature auxospores and initial cells.

### Imaging flow cytometry (iFCM) and confocal imaging

A subsample of each mating culture was fixed in 1% glutaraldehyde at the same time as the loading of the Rhapsody cartridge. Fixed cultures kept at 4°C were subsequently stained with 1% SYBR Green and fed into an ImageStream X Mark II imaging Flow Cytometer (Amnis) for cell density estimation and morphological characterisation of sexual size restoration. Objects in focus were selected using the “Gradient RMS” parameter, followed by filtering for *C. closterium* cells by gating on both the nuclear SYBR Green (excitation laser 488 nm, 0.5 mW) and chloroplast autofluorescence signal (excitation laser 642 nm, 0.5 mW). Microscopic pictures of each cell were visually inspected in the bright field (LED, 13.22 mW) and SYBR Green channels. Occasional cell clumps and other artifacts were flagged for removal. The DNA ploidy level was determined by gating the SYBR Green channel histogram into a diploid (2C) and a post-replication diploid (4C) population. Haploid cells (1C) could not be distinguished from the 2C peak. Next, cells were scored into a cell type category based on their shape and number of nuclei: (1) vegetative cells, (2) binucleate gametangia after meiosis I with visible cleavage furrow, (3) tetranucleate gametangia after meiosis II (4) gametes or zygotes, and (5) expanding auxospores. Vegetative cells were further subdivided based on cell length, with MT- being the larger parental strain. Sexual cells were rare in the iFCM data, making up only 2.2% of all imaged cells (141/6520), possibly because they are more fragile. Doublets of paired gametangia were rarely observed because they likely were separated during cell suspension. To investigate the timing of nuclear fusion, the strains ZW2.20 and CA1.15 were crossed for 25h followed by fixation in glutaraldehyde and staining with 0.3x SYBR Green solution. Developing auxospores were imaged in a covered Lab-Tek II chamber using a Leica TCS SP8X confocal microscope (Leica microsystems) with a 63X (HC PL APO CS2, NA = 1.20) water -corrected objective. For imaging eGFP, a white light laser (WLL) at an excitation wavelength of 488 nm and emission bandwidth of 484-541 nm was used. The autofluorescence signal from the plastids was acquired using a WLL with excitation at 580 nm and emission bandwidth of 645-726 nm. Images were acquired via line sequential imaging on hybrid detectors (HyDTM) in standard mode, without the use of gating.

### Massive parallel single-cell RNA sequencing

A high-throughput procedure for diatom single-cell transcriptomics was established, based on the capture of live algal cells in microwells of the BD Rhapsody Single-Cell Analysis System, precluding the need for protoplasting or nucleus extraction and thus allowing cells to stay in their native medium. Specifically, surface-attached benthic *C. closterium* cells were suspended using a cell scraper and flasks were gently vortexed to separate cells. Next, 575 µL of culture was loaded on the BD Rhapsody cartridge, corresponding to approximately 53,300 (12h), 60,100 (19h) and 98,400 cells (24h). After 15 minutes of sedimentation, large auxospores that exceeded the microwell diameter were forced into the wells through 1 min of centrifugation of the cartridge with a speed of 500 g. Then, cells were lysed with the standard BD lysis bufferfor at least 3 min after which reverse transcription and Exonuclease I treatment were performed according to manufacturer’s instructions. Random priming and extension, cDNA amplification and index PCR were performed following manufacturer’s instructions (23-24117(02)) to obtain a 3’ directed sequencing library. Libraries were sequenced on a NovaSeq 6000 SP flow cell at the Nucleomics core (www.nucleomics.be) with 62-8-0-90 configuration for R1-i7-i5-R2 respectively.

### Mapping, quality filtering, dimensionality reduction, clustering and data integration

STAR v2.5.1b was used to generate the mapping index, after which the BD Rhapsody Sequence Analysis Pipeline (BD; version 1.11) was used to map the FASTQ files to the reference genome. To increase mapping rates, coding sequences in the reference genome were extended with 500bp 3’UTR regions or until the start of the neighboring gene using https://github.com/danilotat/UTR_add_extend_GTF. Single-cell analysis was performed using RStudio (R version 4.2.0) starting from the complete, unfiltered matrix. Cells containing less than 200 genes and genes expressed in less than 2 cells were filtered out of the count matrix.

Outliers were identified based on two metrics: cells containing fewer than 250 or more than 50,000 unique transcripts were filtered out, as were the cells expressing fewer than 250 or more than 9000 genes. The SCTransform function from the Seurat v4 package ^60^ was used to normalize the counts, define the 3000 highly variable genes and scale these features for dimensionality reduction. The first 20 principal components were used to generate UMAPs of individual replicates. Subsequently, the three samples were merged using the merge function from Seurat, after which SCTransform was performed once again. The first 25 principal components were used to generate the merged UMAP plot. Cells were clustered into 16 distinct clusters by the FindClusters function in Seurat, using a resolution of 0.8.

### Identification of trajectory-based and clustering-based marker genes

Clustering-based marker genes were identified using the FindAllMarkers function in Seurat v5.1.0 ^61^ with default settings (Wilcoxon Rank Sum test) but setting the log2 fold change threshold to 0.5. The same settings were used to determine genotype-biased genes among the cells of Cluster 1 and 2. An Euler diagram assessing the overlap between clustering-based and trajectory-based marker genes was created using the eulerr package for R ^62^. All marker genes were annotated with InterPro domains and assigned to homologous gene families from Audoor et al. (2024) ^10^. Enrichment of InterPro domains and gene families in each of the 16 different cell clusters against the full set of 24,359 protein-coding genes as a background was performed using the enricher function of the ClusterProfiler package for R ^63^. We considered features significantly enriched when they had a false discovery rate (FDR) adjusted p-value below 0.01 and comprise at least four marker genes in that cluster.

In parallel, a trajectory spanning the entire dataset was fitted to the UMAP plot using Slingshot v2.12.0 ^64^, specifying the starting (Cluster 1) and final cluster (Cluster 16). The pseudotime for each cell was determined using Slingshot’s slingPseudotime function. For each gene, a negative binomial generalized additive model was fitted that models expression (#UMIs) in function of pseudotime using the figGAM function of the tradeSeq package ^65^, specifying 8 knots. Genes whose expression was significantly associated with pseudotime were detected using the AssociationTest function from tradeSeq. Association was determined over 14 points across the lineage against a log2 fold change cutoff of 1.7 relative to the end of the trajectory (contrastType = “end”). To avoid detection of random variability by extremely lowly expressed genes, we considered only those 8139 genes that are expressed in at least 50 cells with at least 2 UMIs. The significance level for associated genes was controlled on a 5% FDR level using the Benjamini-Hochberg procedure. For visualization, predicted mean smoother values over pseudotime were extracted using the predictSmooth function in tradeSeq for a grid of 500 points. To link pseudotime to real developmental time, we fitted generalized additive models (GAMs) for each cell type from the standardized time series experiment, and determined the time at which cell types reached 5% of the total cell population. In parallel, we projected all scRNA-seq cells to the trajectory and determined the pseudotime where a new cell type starts to dominate (>50%) the dataset. The time intervals between successive cell types in real developmental time and pseudotime were correlated using linear regression.

### Cell type annotation and characterization through reference genes

Known reference genes were used to pinpoint specific cellular events and annotate sexual cell types in the scRNA-seq dataset. A set of six meiosis-specific and three auxospore-associated *C. closterium* genes were retrieved from Audoor et al. 2024 ^10^ and diatom cell wall biosynthesis gene families were taken from Bilcke et al. 2021 ^66^. Genes involved in the cytoskeleton (actin, myosin, tubulin) were identified in *C. closterium* through phylogenetic analysis. Multiple sequence alignments of protein fasta files containing *C. closterium* candidates and reference proteins in other (diatom + non-diatom) species were performed with MAFFT v7.453, followed by automatic trimming of positions containing more than 50% of gaps using trimal v1.4.1 and bootstrap consensus tree generation using IQ-tree v2.2.2.6 with 1000 ultrafast bootstrap repeats ^67^. Finally, curated sets of *C. closterium* reference genes for the S-phase (n = 32) and M-phase (n = 7) were used to determine cell cycle stages using the CellCycleScoring functionality in Seurat v5.1.0 ^61^. Protein structure predictions were generated with Alphafold 3 ^68^ and visualized with PyMOL 3.0.0. Transmembrane domains were predicted with DeepTMHMM ^69^ and signal peptides with Phobius (https://phobius.sbc.su.se/).

To identify the ancient karyogamy gene *GEX1* in diatoms, an initial set of known GEX1 protein sequences in plants, yeast, and the SAR clade (Rhizaria, Alveolates and non-diatom Stramenopiles) ^2^ were aligned with MAFFT v7.453 and a profile Hidden Markov Model (HMM) was created using the hmmbuild command of hmmer v3.1b2. The hmmsearch command was used to identify distant GEX1 homologs (E < 1e-4) in the PLAZA Diatoms protein database ^48^ and *C. closterium* ^10^ and *S. marinoi* ^70^ proteomes. The resulting diatom GEX1 sequences were extracted for a reverse HMM search against the PLAZA Diatoms database, confirming that the closest homologs (E < 1e-4) are *A. thaliana* and *E. siliculosis* GEX1 proteins. For phylogenetic analysis, GEX1 protein sequences from various databases (NCBI Protein, the Marine Microbial Eukaryote Transcriptome Sequencing Project (MMETSP) ^71^ and PLAZA Diatoms) for different eukaryotic clades (animals, fungi, Viridiplantae, SAR clade) were aligned with MAFFT v7.453 and a bootstrap consensus tree was generated using IQ-tree v2.2.2.6 with 1000 ultrafast bootstrap repeats ^67^. Because of low sequence conservation, no trimming was performed.

Expression of genes of interest was visualized in UMAP plots using the FeaturePlot function from Seuratv5.1.0 ^61^. In case of multiple reference genes for the same process, the AddModuleScore_UCell function from the UCell package ^72^ was used to calculate signature enrichment for the reference gene set through the Mann-Whitney U statistic.

### Cell type annotation through bulk RNA-seq

The expression pattern of single cells was cross-referenced with differential expression results from a bulk RNA-seq experiment that contrasts sexual versus control expression across three time points (T1 = 9h, T2 = 14h and T3 = 27h) ^10^. Four classes of indicator genes were defined based on their expression pattern in bulk RNA-seq: “Down indicators”: union of the 100 top most significantly downregulated genes during sex in each of the three time points, and “T1,T2,T3 indicators”: the 100 most significantly upregulated genes in T1, T2 and T3 respectively whose fold changes exceeded that of the other timepoints. For each cell, the summed UMI counts for each indicator class were calculated and the relative contribution of each indicator to the cellular transcriptome was visualized using pie charts with the scatterpie package v0.2.3 for ggplot2. In addition, the response of ZW2.20 MT+ cultures to a filtrate containing sex-inducing pheromone minus (SIP-) was assessed using bulk RNA-seq of matched case-control samples at two time points after addition of the pheromone (3h, 9h), as described in Belišová et al. 2024 ^73^.

### Distinguishing cells by genotype and mating type

Genotyping of the MT+ strain ZW2.20 was performed by harvesting 100 mL of cell culture in the late exponential phase by filtration on a Versapor filter (3 µm pore size, 25 mm diameter, PALL). Next, filters with cell material were flash-frozen in Eppendorf tubes and stored at -80°C until DNA extraction. High-molecular-weight DNA for whole-genome sequencing was extracted using an optimised CTAB protocol ^10^. The DNA concentration and quality were evaluated by spectrophotometry (Nanodrop, Thermo Scientific) and agarose gel electrophoresis. Library preparation was performed using the NEBNext® Ultra™ II DNA Library Prep Kit and a total of 56,233,526 paired-end 150bp Illumina reads were sequenced at the Nucleomics core (www.nucleomics.be). Read quality control was performed with FastQC version 0.11.2 ^74^. ZW2.20 reads were mapped to the CA1.15 reference genome using bwa v0.7.17 ^75^ and duplicates were tagged with Samblaster v0.1.26 ^76^. Conversion to bam, sorting and indexing was performed with samtools v1.15.1 ^77^. Variant calling was performed using bcftools 1.15.1, using the mpileup function to generate genotype likelihoods, followed by variant calling using the settings -m -O v -g 8. Next, the bcftools “filter” function was used to remove variants within a minimum SNP gap of 3 (-g3), an indel gap of 10 (-G10), had a quality score below 10, a combined depth under 5 or maximum depth over 500. This resulted in a final filtered variant set of 394,873 single-nucleotide polymorphisms and 50,717 indels.

In parallel, single-cells were demultiplexed into genotypes using Souporcell ^78^. Specifically, the aligned reads contained in the BD analysis pipeline BAM-file were used as an input into the Souporcell algorithm with k = 2 to cluster cells into two distinct populations and a recombined “doublet” population. Afterwards, Demuxafy was used to assign the correct identity to each identified population based on the ZW2.20 versus CA1.15 variant signature (VCF file) ^79^.

The mating type identity of single cells was independently determined by the expression of genes with a mating type-specific expression in vegetative (non-sexual) cells, as defined in Belisova et al. (2024) ^73^. In brief, we used all 75 MT+ and 11 MT- biased genes that were upregulated in large and small cells or only in small cells. The bias of single-cells towards either mating type was determined with the AddModuleScore function from Seurat v5.1.0 ^61^. The mating type with the highest module score was selected, except when both scores were negative, then the outcome was deemed undetermined.

### Identification of genotype-specific genes

To compare the expression of different genotypes across pseudotime, we focused on cells that were assigned to a single genotype (no doublets/offspring) and fell along the differentiated mating type-specific trajectories by keeping 2570 CA1.15 (MT-) and 2693 ZW2.20 (MT+) cells (**Fig. 3e**). Strongly differentially expressed genes between these two populations were determined using the “FindAllMarkers” function of Seurat using a log2 fold change cutoff of 3. To verify whether genes appear MT- specific because of a failure of ZW2.20 to map against the CA1.15 genome, we used the intersect function of bedtools v2.2.28 to subset the previously generated VCF file per gene and determined the number of mismatches (by SNPs or indels). Genes with a mating type-specific expression (median: 13 mismatches) were not more diverged than all other genes in the genome (median: 14 mismatches), suggesting that mating type-specific expression is, in general, not caused by genomic divergence. Trajectories were fitted through the parallel ZW2.20 and CA1.15 populations with Slingshot v2.12.0 ^64^. The pseudotime for each cell was determined using Slingshot’s slingPseudotime function, after which a pseudotime offset of 3.5 was added for the MT- trajectory to synchronize the trajectories. Next, smooth GAMs with 8 knots were fitted using TradeSeq v1.8.0 to model expression along the mating type-specific trajectories. The predicted expression of GAMs at 60 discrete points along pseudotime was converted to distance, defined as the complement of the Pearson correlation (1-cor). Then, GAMs with similar expression dynamics were hierarchically clustered into modules using the hclust command in R and the resulting dendrograms were visualized using ggdendroplot. Functional clustering of genotype-biased genes on the *C. closterium* genome was determined using C-hunter v1.0 ^80^ based on a custom directed acyclic graph ontology representing both the differences in genotypes and pseudotime module (four for ZW2.20, four for CA1.15) and setting the minimum cluster size to 3 genes. Functional clustering on the 100 longest contigs was visualized using the Circlize v0.4.16 package for R ^81^.

### Experimental validation using transgenic reporter lines

Vector assembly was based on the Greengate cloning system ^82^. Sequences for cloning into the entry modules were obtained through PCR amplification of the target sequence with a 20 bp overlap with the entry module. In this way, the promoter regions of AAE2 (Ccl_15126, 603 bp), GEX1 (Ccl_2478, 564 bp) and two MYB3R5 homologs (Ccl_1772, 475 bp and Ccl_1778, 363 bp) were cloned from the end of the previous gene to the translation start site. The promoter was then fused with enhanced GFP (eGFP) and the *Cylindrotheca fusiformis* fcp terminator in a high-copy number pGG-AG-KmR destination vector ^83^. To introduce pAAE2:eGFP into strain CA1.15 (MT-), we used biolistic transformation using a cotransformation approach with a plasmid carrying a nourseothricin resistance (NrsR) cassette controlled by the *C. fusiformis* fcp promotor and terminator (pCf_fcp:NrsR). In brief, 3 mg of gold particles (0.6 µm diameter, Bio-Rad) were coated with 5 µg of reporter plasmid and 5 µg of selection plasmid using CaCl2 and spermidine according to the manufacturer’s instructions. Particles were accelerated towards one hundred million (1 x 10⁸) agar-plated *C. closterium* cells at a pressure of 1550 psi using the PDS-1000/He System (Bio-rad). After 24h, cells were transferred to 1.5% ASW agar plates containing 200 μg/mL clonNAT (nourseothricin sulfate). Identical steps were performed for pCcl_1772:eGFP and pCcl_1778:eGFP, but instead using the NrsR resistance cassette cloned on the same vector. On the other hand, pGEX1:eGFP was introduced into the MT- using bacterial conjugation with DH5α pTA-Mob *Escherichia coli*, as described by *Karas et al. 2015* ^84^. Here, the nourseothricin resistance cassette was cloned on the same vector as pGEX1:eGFP, together with a centromere autonomous replicating sequence CEN6-ARS4-HIS3 (CAH). After 48h on LB+ 1% ASW agar plates, the cells were transferred to 1.5% ASW agar plates containing 200 ug/mL clonNAT.

Colonies were picked after three weeks of selection and transferred to liquid medium containing 200 μg/mL clonNAT. After 10 days, 500 μL subcultures of these mutant colonies were transferred to eppendorf tubes and harvested by centrifugation (1 min, 10.000 rpm). DNA was extracted using a lysis buffer (1% TritonX-100, 20 mM Tris-HCl, pH8, 2mM EDTA) followed by vortexing, cooling on ice and a heating step at 85 degrees for 10 minutes. With the resulting DNA as a template, genotyping for enhanced green fluorescent protein (eGFP) by PCR was used to to verify the presence of the introduced plasmid.

To pinpoint gene expression during sexual development, transgenic lines were crossed with a wild type strain of the opposite mating type. Cultures were inoculated at 200 cells/mL in small 25 cm² cell culture flasks (VWR®) filled with 20 mL of artificial seawater. Four days later, cultures were subjected to a 36h dark arrest after which mating was initiated, still in the dark, by pooling 50 µL of each mating type in a covered Lab-Tek II chamber with ASW. After crossing, the chambers were put in continuous light and imaged using a Leica TCS SP8X confocal microscope (Leica microsystems) with a 63X (HC PL APO CS2, NA = 1.20) water immersion-corrected objective at different time points after crossing (6h, 21h, 24h, 25h, 26h and 40h). Images were acquired with hybrid detectors (HyDTM) in standard mode using a time gated window between 1 and 12 ns, in a line sequential mode. For imaging eGFP expression, a white light laser (WLL) was used at an excitation wavelength of 488 nm and emission bandwidth of 494-543 nm. The autofluorescence of plastids was acquired using a WLL with excitation at 580 nm and emission bandwidth of 625-663 nm. To confirm the absence of expression in non-sexual cells, non-crossed transgenic cultures were included for imaging as a negative control. The cells were imaged as described above at different timepoints after the cross.

### Annotation of transcription factors and orthology-based binding sites

*C. closterium* TFs were defined by the presence of at least one DNA-binding InterPro domain for a single type of TF (referred to here as a “TF family”), leading to the identification of 481 putative TFs. An additional 15 Myb and C2H2 TFs that were previously annotated by Rayko in *P. tricornutum* ^43^ were identified in *C. closterium* using the orthology method from Audoor et al. ^10^. The putative TFs that were significantly associated with pseudotime during sex were manually curated to retain only those genes belonging to TF families with unambiguous TF activity. The *C. closterium* Myb family was annotated using phylogenetic analysis with MAFFT and IQ-tree as explained above, based on reference Myb and Myb-like proteins in *P. tricornutum* and *T. pseudonana* from Rayko et al. ^43^ and Wang et al. ^85^. Differential expression data was plotted onto the Myb family tree using ggtree ^86^.

An orthology-based approach was used to identify known eukaryotic TF binding sites in the promoter region of all *C. closterium* genes. First, we extracted the promoter region upstream of the translation start site, taking a 500-bp region or up to the previous gene model. We selected the translation start site because important cis-regulatory elements can also be located in the untranslated regions ^87^. Then, for each predicted *C. closterium* TF, we identified potential orthologs in human (*Homo sapiens*), *Arabidopsis thaliana* and yeast (*Saccharomyces cerevisiae)*, selecting those TFs with the most PLAZA orthology evidences (Tree-based ortholog, orthologous gene family and Best-Hits-and-Inparalogs family) using the integrative orthology dataset from Audoor et al. ^10^. This led to the identification of one or more orthologs for 315 *C. closterium* TFs. Motifs associated with the orthologous TFs were extracted from CisBP ^88^, enriched with additional motifs from JASPAR 2022 ^89^ and Kulkarni et al. 2019 ^90^ for *A. thaliana*. The orthology-based motifs per *C. closterium* TF were then gathered and mapped to the promoter region using FIMO v5.5.5 ^91^ using the default p-value cutoff of 1e-4 and inferring the background model from the promoter fasta file, and Cluster-Buster v2017-09-22 ^92^ with a cluster score threshold of 5, a motif score threshold of 6 and an expected gap between neighbouring motifs of 35. This resulted in a feature file that defines which motifs were mapped to the promoter of each gene, either by FIMO or Cluster-Buster.

### Machine learning models for the identification of *de novo* transcription factor binding sites

Clustering-based marker genes were assigned to five broader cell types: vegetative, mate finding, paired gametangia, gametes and auxospores (**Figure 2c**). Markers for multiple cell types were assigned to the cell type where they were most significantly upregulated. This resulted in a unique set of markers for each cell type, which was used as a positive class to train machine learning (ML) models. Meanwhile, we defined non-responsive genes as a control class for binary classification: genes that were not a significant marker for any cluster, and had the most stable expression across the entire dataset, as defined by the coefficient of variation over the average UMI counts of all clusters. Next, undersampling was performed by binning non-responsive genes in ten quantiles by expression level, and selecting the most stable genes (lowest coefficient of variation) from each bin to achieve a similar number of non-responsive and marker genes for each cell type.

Important *de novo* motifs for each cell type were identified in the 500-bp promoter upstream of the translation start site using two general strategies: (i) enrichment analysis of markers versus non-responsive genes with the oligo-diff command from RSAT ^93^ for k-mers of 5-9 bp as well as with STREME v5.5.5 ^94^ for 6-12 bp k-mers, and (ii) identification of overrepresented sequences in markers’ promoters using cisDIVERSITY version 2.8.2 ^95^ for PWMs (position weight matrices) with a minimum length of 6. The most likely transcription factor family that binds each motif was identified by selecting the most significant hit from a TomTom search (-no-ssc -verbosity 1 -min-overlap 5 -dist pearson -evalue -thresh 10.0 <top features>) against the eukaryotic non-redundant JASPAR 2022 database. We then mapped these *de novo* motifs back to the 500-bp promoters of *C. closterium* genes using FIMO and Cluster-Buster as explained above, generating a feature file with the maximum number of occurrences of each motif in each promoter.

For each cell type, we built random forest machine learning models that predict cell type marker identity based on the number of occurrences of motifs in a gene’s promoter using the method of Smet and colleagues ^96^. Specifically, Random Forest ML models were trained using scikit-learn for Python ^97^ with automatic hyperparameter optimization using Optuna ^98^ to maximize the F1 score during a five-fold cross-validation, followed by training of the classifier with optimal hyperparameters on the entire training set (75% of data) and evaluation of performance on the test set (25%). For each cell type, ten different models were trained with random train-test splits. Afterwards, a SHAP (SHapley Additive exPlanations) analysis was performed for each model to determine the contribution of each motif to the model’s predictions ^99^. The top-5 most important motifs in each of these ten models were merged and selected for construction of gene regulatory networks.

### Generation of gene regulatory networks

An omnibus set of all predicted motifs in the 500-bp promoters of *C. closterium* genes was created by merging orthology-based motifs with machine learning-based motifs. A gene regulatory network was inferred with MNI-EX v1 ^100^ using default settings, including the expansion of motifs to the TF family level for enrichment but skipping the functional enrichment stage. For each regulon, hypergeometric tests were performed with the enricher function of the ClusterProfiler package for R ^63^ to determine whether target genes are enriched in specific interpro domains or stages of the cell cycle. Gene regulatory networks were visualized with Cytoscape 3.10.2 ^101^.

### Comparative (meta)transcriptomic analyses

For comparative transcriptomics analysis of GEX1 and Myb transcription factors, bulk RNA-seq expression values (TMM normalized counts per million), log2 fold changes and differential expression statistics in four diatom species (*C. closterium, Seminavis robusta, P. multistriata, S. marinoi*) were retrieved from Audoor et al. 2024 ^10^. Myb TFs in the latter three species were identified as the union of all genes that carry Myb InterPro domains as well as those belonging to the previously described Myb subfamily HOM338GF000037 ^10^. These Mybs were classified by adding them to the three-species phylogenetic framework of Mybs described above.

Protein sequences, expression data, differential expression calls and Kraken2 taxonomic labels from a salinity-treated metatranscriptome experiment from the Scheldt estuary in Belgium were retrieved from Bilcke et al. (2025) ^46^. Protein fasta files were filtered for Thalassiosirales sequences (143,872 proteins) and were annotated with InterProScan v5.75 using the following models: CDD, Gene3D, PRINTS, SUPERFAMILY, SMART, PIRSF, Pfam ^102^. The environmental Myb TFs with the highest upregulation (log2FC > 5) were phylogenetically classified in the Myb family as explained above. In particular, Rayko et al.’s *T. pseudonana* tp_Myb2R2 ^43^ was positioned together with pennate Myb2R1 homologs rather than with the pennate Myb2R2, making it hard to classify (environmental) Myb2R2 homologs from centrics with certainty. A set of six sex-specific Myb3R5 homologs in the Scheldt microcosm metatranscriptome was further taxonomically annotated within the Thalassiosirales by constructing a bootstrap-consensus phylogenetic tree with IQ-tree that includes reference Thalassiosirales Myb3R5 sequences from Roberts et al. 2023 ^103^.

Homologs for Myb3R5 were identified in the proteomes predicted from *Tara* Oceans metagenome-assembled genomes (MAGs), downloaded from www.genoscope.cns.fr/tara/ (file path: www.genoscope.cns.fr/tara/localdata/data/SMAGs-v1/SMAGs_v1_concat.faa.tar.gz, downloaded on 16/01/2026). Specifically, a HMM search for Myb3R5 was performed against the protein sequences encoded on 52 diatom MAGs (E-value < 1e-50). Next, an IQ-tree phylogenetic tree was constructed as explained above, combining HMM hits and reference protein sequences for the entire Myb family, followed by classification of 36 *bona fide* Myb3R5 *Tara* transcripts from 26 diatom MAGs. In parallel, we retrieved expression data mapped to the genes of 27 meiotic-positive (expressing SPO11-2 at least ones) diatom MAGs in *Tara* Oceans and *Tara Polar Circle* from Bilcke et al. 2025 ^46^ (https://doi.org/10.5281/zenodo.11258438 - Tara_MAG_coexpression.zip - MAG_level_TPM_DiatomMAGs.txt). In this dataset, expression was normalized for genotype abundance by calculating transcripts per million (TPM) on the MAG level. Next, we filtered this dataset for those MAGs (n = 17) that express Myb3R5 with at least 4 reads. The MAG-level TPM expression of filtered Myb3R5 genes was visualized on a world map by combining the “ggplot2” and “maps” packages for R, using coordinates for each *Tara* station downloaded from https://zenodo.org/records/16637330. Finally, we correlated the Myb3R5 expression level (TPM) with the sexual signal within each MAG in each sample, by determining the number of previously described sex markers (pennate: 4; centric: 5 markers) that are co-expressed relative to a predefined TPM threshold ^46^. Log transformed Myb3R5 expression was modelled in function of the discrete variable “number of sex markers”, using a linear model (ANOVA) and t-tests comparing each group with the intercept (0 markers co-expressed).

## Author contributions

GB, LDV, WV and KV conceived of the study and directed research. GB, NR, TE, CG and BDR developed the protocol for single-cell transcriptomics in diatoms. Sexual crosses for scRNA-seq were performed by NR. Imaging flow cytometry and subsequent visual annotation of cell types was performed by PC, NR and DB. AC, SA and DB established biolistic and conjugation-based transformation of *Cylindrotheca closterium*. AC constructed all transcriptional reporter lines with help of JP. Confocal imaging was performed by AC with input from GB, EM, DB, PC and DVD. SA performed the standardized cross of *C. closterium* for chronology estimation. DB sequenced C*. closterium* strain ZW2.20 for genotyping. Transcription factor identification, and the motif identification and ranking with machine learning were performed by TF with contributions by NMP. Single-cell transcriptomic, comparative genomic, phylogenomic and metatranscriptomic analyses were carried out by GB, NR and TE. MVB applied the functional clustering analysis. The manuscript was initially written by GB, LDV, WV and KV based on input from AC, NR, TF and BDR, and all authors contributed to the final text.

## Acknowledgements

We thank Thomas Depuydt, Petra Bulankova, Koen Van den Berge, Mariella Ferrante, Antonella Ruggiero, Tobias Gerber, Dajo Smet, Svitlana Lukicheva and Koen Sabbe for helpful discussion about our dataset.

## Resource availability

Raw scRNA-seq Illumina reads, unfiltered and filtered count matrices and the merged Seurat object containing all relevant metadata (cell type, sample, clustering, genotype, cell cycle phasing, …) is available in NCBI GEO with accession number GSE303315. Whole-genome Illumina sequencing libraries of strain ZW2.20 for genotyping are available at the European Nucleotide Archive with accession PRJEB83002. Tables with marker genes and their functional annotation, gene regulatory network files, as well as details about vector assembly are available on Zenodo with DOI: https://doi.org/10.5281/zenodo.15863610. Raw paired-end Illumina reads of the sex-inducing pheromone experiment can be found at the European Nucleotide Archive (PRJEB81513). Transgenic *C. closterium* cultures (reporter lines) were deposited to the BCCM/DCG culture collection for cryopreservation: DCG 1335 (AAE2), DCG 1388 (Ccl_1772, Myb3R5), DCG 1389 (Ccl_1778, Myb3R5) and DCG 1390 (GEX1). *Cylindrotheca closterium* CA1.15 gene models are available at bioinformatics.psb.ugent.be/gdb/Cylindrotheca_closterium/Version1.2/

## Bibliography

1. Vosseberg, J. et al. The emerging view on the origin and early evolution of eukaryotic cells. Nature 633, 295–305 (2024).

2. Speijer, D., Lukeš, J. & Eliáš, M. Sex is a ubiquitous, ancient, and inherent attribute of eukaryotic life. Preprint at 10.1073/pnas.1501725112 (2015).

3. Dunthorn, M. & Katz, L. A. Secretive ciliates and putative asexuality in microbial eukaryotes. Trends in Microbiology 18, 183–188 (2010).

4. Rizos, I., Frada, M. J., Bittner, L. & Not, F. Life cycle strategies in free-living unicellular eukaryotes: Diversity, evolution, and current molecular tools to unravel the private life of microorganisms. Journal of Eukaryotic Microbiology 71, e13052 (2024).

5. Field, C. B., Behrenfeld, M. J., Randerson, J. T. & Falkowski, P. Primary production of the biosphere: Integrating terrestrial and oceanic components. Science 281, 237–240 (1998).

6. Nakov, T., Beaulieu, J. M. & Alverson, A. J. Accelerated diversification is related to life history and locomotion in a hyperdiverse lineage of microbial eukaryotes (Diatoms, Bacillariophyta). New Phytologist 219, 462–473 (2018).

7. Alverson, A. J. et al. Phylogenomics reveals the slow-burning fuse of diatom evolution. Proceedings of the National Academy of Sciences 122, e2500153122 (2025).

8. Davidovich, N. A. et al. Ardissonea crystallina has a type of sexual reproduction that is unusual for centric diatoms. Sci Rep 7, 14670 (2017).

9. Kaczmarska, I. et al. Proposals for a terminology for diatom sexual reproduction, auxospores and resting stages. Diatom Research 28, 263–294 (2013).

10. Audoor, S. et al. Transcriptional chronology reveals conserved genes involved in pennate diatom sexual reproduction. Mol Ecol 33, e17320 (2024).

11. Bilcke, G. et al. Mating type specific transcriptomic response to sex inducing pheromone in the pennate diatom *Seminavis robusta*. ISME Journal 15, 562–576 (2021).

12. Basu, S. et al. Finding a partner in the ocean: molecular and evolutionary bases of the response to sexual cues in a planktonic diatom. New Phytologist 215, 140–156 (2017).

13. Annunziata, R. et al. Trade-off between sex and growth in diatoms: Molecular mechanisms and demographic implications. Science Advances 8, eabj9466 (2022).

14. Grujčić, V. et al. Towards high-throughput parallel imaging and single-cell transcriptomics of microbial eukaryotic plankton. PLOS ONE 19, e0296672 (2024).

15. Ku, C. et al. A single-cell view on alga-virus interactions reveals sequential transcriptional programs and infection states. Science Advances 6, eaba4137 (2020).

16. Fromm, A. et al. Single-cell RNA-seq of the rare virosphere reveals the native hosts of giant viruses in the marine environment. Nat Microbiol 9, 1619–1629 (2024).

17. Hevroni, G., Vincent, F., Ku, C., Sheyn, U. & Vardi, A. Daily turnover of active giant virus infection during algal blooms revealed by single-cell transcriptomics. Science Advances 9, eadf7971 (2023).

18. Gatt, C., Xie, Y., Wahi, K., Johansson, E. M. & Zanini, F. Integrating microscopy and transcriptomics from individual uncultured eukaryotic plankton. eLife 13, (2024).

19. Ma, F., Salomé, P. A., Merchant, S. S. & Pellegrini, M. Single-Cell RNA sequencing of batch Chlamydomonas cultures reveals heterogeneity in their diurnal cycle phase. The Plant Cell koab025 (2021) doi:10.1093/plcell/koab025.

20. Belišová, D. et al. Molecular fingerprints of cell size sensing and mating type differentiation in pennate diatoms. New Phytologist 245, 1625–1639 (2025).

21. Jiang, Y., Adhikari, D., Li, C. & Zhou, X. Spatiotemporal regulation of maternal mRNAs during vertebrate oocyte meiotic maturation. Biological Reviews 98, 900–930 (2023).

22. Carrell, D. T. Epigenetics of the male gamete. Fertil Steril 97, 267–274 (2012).

23. Rutley, N. & Twell, D. A decade of pollen transcriptomics. Plant Reprod 28, 73–89 (2015).

24. Lee, U., Li, C., Langer, C. B., Svetec, N. & Zhao, L. Comparative single-cell analysis of transcriptional bursting reveals the role of genome organization in de novo transcript origination. Proceedings of the National Academy of Sciences 122, e2425618122 (2025).

25. Geijer, C. et al. Time course gene expression profiling of yeast spore germination reveals a network of transcription factors orchestrating the global response. BMC Genomics 13, 554 (2012).

26. White, R. J. et al. A high-resolution mRNA expression time course of embryonic development in zebrafish. eLife 6, e30860.

27. Davutoglu, M. G. et al. Gliding motility of the diatom Craspedostauros australis coincides with the intracellular movement of raphid-specific myosins. Commun Biol 7, 1–13 (2024).

28. Vanstechelman, I., Sabbe, K., Vyverman, W., Vanormelingen, P. & Vuylsteke, M. Linkage mapping identifies the sex determining region as a single locus in the pennate diatom Seminavis robusta. PLoS ONE 8, e60132 (2013).

29. Russo, M. T. et al. MRP3 is a sex determining gene in the diatom Pseudo-nitzschia multistriata. Nature Communications 9, 5050 (2018).

30. Bolwell, G. P., Callow, J. A. & Evans, L. V. Fertilization in brown algae. III. Preliminary characterization of putative gamete receptors from eggs and sperm of Fucus serratus. J Cell Sci 43, 209–224 (1980).

31. Catt, J. W., Vithanage, H. I. M. V., Callow, J. A., Callow, M. E. & Evans, L. V. Fertilization in brown algae: V. Further investigations of lectins as surface probes. Experimental Cell Research 147, 127–133 (1983).

32. Callow, J. A. Sexual recognition and fertilization in brown algae. J Cell Sci Suppl 2, 219–232 (1985).

33. Baroux, C., Autran, D., Gillmor, C. S., Grimanelli, D. & Grossniklaus, U. The maternal to zygotic transition in animals and plants. Cold Spring Harb Symp Quant Biol 73, 89–100 (2008).

34. Zhao, P. et al. Two-Step Maternal-to-Zygotic Transition with Two-Phase Parental Genome Contributions. Dev Cell 49, 882–893.e5 (2019).

35. Ferrante, M. I. et al. Exploring molecular signs of sex in the marine diatom *Skeletonema marinoi*. Genes 10, 494 (2019).

36. Alandete-Saez, M., Ron, M., Leiboff, S. & McCormick, S. Arabidopsis thaliana GEX1 has dual functions in gametophyte development and early embryogenesis. The Plant Journal 68, 620–632 (2011).

37. Nishikawa, S. I. et al. Arabidopsis GEX1 is a nuclear membrane protein of gametes required for nuclear fusion during reproduction. Frontiers in Plant Science 11, 548032 (2020).

38. Bogaert, K. A., Beeckman, T. & De Clerck, O. Egg activation-triggered shape change in the Dictyota dichotoma (Phaeophyceae) zygote is actin-myosin and secretion dependent. Ann Bot 120, 529–538 (2017).

39. Schreiber, V. et al. The Central Vacuole of the Diatom Phaeodactylum tricornutum: Identification of New Vacuolar Membrane Proteins and of a Functional Di-leucine-based Targeting Motif. Protist 168, 271–282 (2017).

40. Davidovich, N. A. Transition to sexual reproduction and control of initial cell size in *Nitzschia lanceolata*. Diatom Research 13, 29–38 (1998).

41. Vanormelingen, P. et al. Heterothallic sexual reproduction in the model diatom Cylindrotheca. European Journal of Phycology 48, 93–105 (2013).

42. Schmid, A.-M. M. Aspects of morphogenesis and function of diatom cell walls with implications for taxonomy. Protoplasma 181, 43–60 (1994).

43. Rayko, E., Maumus, F., Maheswari, U., Jabbari, K. & Bowler, C. Transcription factor families inferred from genome sequences of photosynthetic stramenopiles. New Phytologist 188, 52–66 (2010).

44. Ito, M. Conservation and diversification of three-repeat Myb transcription factors in plants. J Plant Res 118, 61–69 (2005).

45. Okada, M., Akimaru, H., Hou, D.-X., Takahashi, T. & Ishii, S. Myb controls G(2)/M progression by inducing cyclin B expression in the Drosophila eye imaginal disc. EMBO J 21, 675–684 (2002).

46. Bilcke, G. et al. Conserved genetic markers reveal widespread diatom sexual reproduction in the global ocean. Nature Communications 16, 10029 (2025).

47. Malviya, S. et al. Insights into global diatom distribution and diversity in the world’s ocean. Proceedings of the National Academy of Sciences of the United States of America 113, E1516–E1525 (2016).

48. Osuna-Cruz, C. M. et al. The *Seminavis robusta* genome provides insights into the evolutionary adaptations of benthic diatoms. Nature Communications 11, 3320 (2020).

49. von Dassow, P. & Montresor, M. Unveiling the mysteries of phytoplankton life cycles: patterns and opportunities behind complexity. J Plankton Res 33, 3–12 (2011).

50. Wang, M. et al. Single-Cell RNA Sequencing Analysis Reveals Sequential Cell Fate Transition during Human Spermatogenesis. Cell Stem Cell 23, 599–614.e4 (2018).

51. Wagner, M. et al. Single-cell analysis of human ovarian cortex identifies distinct cell populations but no oogonial stem cells. Nat Commun 11, 1147 (2020).

52. Hou, Z. et al. High-throughput single-cell transcriptomics reveals the female germline differentiation trajectory in Arabidopsis thaliana. Commun Biol 4, 1–16 (2021).

53. Ichino, L., et al. Single-nucleus RNA-seq reveals that MBD5, MBD6, and SILENZIO maintain silencing in the vegetative cell of developing pollen. Cell Reports 41, (2022).

54. Dogga, S. K. et al. A single cell atlas of sexual development in Plasmodium falciparum. Science 384, eadj4088 (2024).

55. Howick, V. M. et al. The Malaria Cell Atlas: Single parasite transcriptomes across the complete Plasmodium life cycle. Science 365, eaaw2619 (2019).

56. Real, E. et al. A single-cell atlas of Plasmodium falciparum transmission through the mosquito. Nat Commun 12, 3196 (2021).

57. Gibson, A. K., Delph, L. F. & Lively, C. M. The two-fold cost of sex: Experimental evidence from a natural system. Evol Lett 1, 6–15 (2017).

58. Mann, D. G. & Vanormelingen, P. An inordinate fondness? the number, distributions, and origins of diatom species. Journal of Eukaryotic Microbiology 60, 414–20 (2013).

59. Behrenfeld, M. J. et al. Thoughts on the evolution and ecological niche of diatoms. Ecological Monographs 91, e01457 (2021).

60. Hao, Y. et al. Integrated analysis of multimodal single-cell data. Cell 184, 3573–3587.e29 (2021).

61. Hao, Y. et al. Dictionary learning for integrative, multimodal and scalable single-cell analysis. Nat Biotechnol 42, 293–304 (2024).

62. Larsson, J., et al. Eulerr: Area-Proportional Euler and Venn Diagrams with Ellipses. R Package Version 6.1.0. (2020).

63. Wu, T. et al. clusterProfiler 4.0: A universal enrichment tool for interpreting omics data. Innovation (Camb) 2, 100141 (2021).

64. Street, K. et al. Slingshot: cell lineage and pseudotime inference for single-cell transcriptomics. BMC Genomics 19, 477 (2018).

65. Van den Berge, K., et al. Trajectory-based differential expression analysis for single-cell sequencing data. Nat Commun 11, 1201 (2020).

66. Bilcke, G. et al. Diurnal transcript profiling of the diatom *Seminavis robusta* reveals adaptations to a benthic lifestyle. The Plant Journal tpj.15291 (2021).

67. Nguyen, L. T., Schmidt, H. A., Von Haeseler, A. & Minh, B. Q. IQ-TREE: A fast and effective stochastic algorithm for estimating maximum-likelihood phylogenies. Molecular Biology and Evolution 32, 268–274 (2015).

68. Abramson, J. et al. Accurate structure prediction of biomolecular interactions with AlphaFold 3. Nature 630, 493–500 (2024).

69. DeepTMHMM predicts alpha and beta transmembrane proteins using deep neural networks | bioRxiv. https://www.biorxiv.org/content/10.1101/2022.04.08.487609v1.

70. Pinseel, E. et al. Strain-specific transcriptional responses overshadow salinity effects in a marine diatom sampled along the Baltic Sea salinity cline. ISME J 16, 1776–1787 (2022).

71. Keeling, P. J. et al. The Marine Microbial Eukaryote Transcriptome Sequencing Project (MMETSP): illuminating the functional diversity of eukaryotic life in the oceans through transcriptome sequencing. PLoS Biology 12, e1001889 (2014).

72. Andreatta, M. & Carmona, S. J. UCell: Robust and scalable single-cell gene signature scoring. Computational and Structural Biotechnology Journal 19, 3796–3798 (2021).

73. Belišová, D. et al. Molecular fingerprints of cell size sensing and mating type differentiation in pennate diatoms. New Phytologist **n/a**,.

74. Andrews, S. FastQC: a quality control tool for high throughput sequence data. Preprint at http://www.bioinformatics.babraham.ac.uk/projects/fastqc/ (2010).

75. Li, H. & Durbin, R. Fast and accurate short read alignment with Burrows–Wheeler transform. Bioinformatics 25, 1754–1760 (2009).

76. Faust, G. G. & Hall, I. M. SAMBLASTER: fast duplicate marking and structural variant read extraction. Bioinformatics 30, 2503–2505 (2014).

77. Li, H. et al. The Sequence Alignment/Map format and SAMtools. Bioinformatics 25, 2078–2079 (2009).

78. Heaton, H. et al. Souporcell: robust clustering of single-cell RNA-seq data by genotype without reference genotypes. Nat Methods 17, 615–620 (2020).

79. Neavin, D. et al. Demuxafy: improvement in droplet assignment by integrating multiple single-cell demultiplexing and doublet detection methods. Genome Biology 25, 94 (2024).

80. Yi, G., Sze, S.-H. & Thon, M. R. Identifying clusters of functionally related genes in genomes. Bioinformatics 23, 1053–1060 (2007).

81. Gu, Z., Gu, L., Eils, R., Schlesner, M. & Brors, B. circlize implements and enhances circular visualization in R. Bioinformatics 30, 2811–2812 (2014).

82. Lampropoulos, A. et al. GreenGate - A Novel, Versatile, and Efficient Cloning System for Plant Transgenesis. PLOS ONE 8, e83043 (2013).

83. Blomme, J., Liu, X., Jacobs, T. B. & De Clerck, O. A molecular toolkit for the green seaweed Ulva mutabilis. Plant Physiol 186, 1442–1454 (2021).

84. Karas, B. J. et al. Designer diatom episomes delivered by bacterial conjugation. Nature Communications 6, 6925 (2015).

85. Wang, W. et al. MYB gene family in the diatom Phaeodactylum tricornutum revealing their potential functions in the adaption to nitrogen deficiency and diurnal cycle. J Phycol 58, 121–132 (2022).

86. Yu, G., Smith, D. K., Zhu, H., Guan, Y. & Lam, T. T.-Y. ggtree: an r package for visualization and annotation of phylogenetic trees with their covariates and other associated data. Methods in Ecology and Evolution 8, 28–36 (2017).

87. Heyndrickx, K. S., Van de Velde, J., Wang, C., Weigel, D. & Vandepoele, K. A functional and evolutionary perspective on transcription factor binding in Arabidopsis thaliana. Plant Cell 26, 3894–3910 (2014).

88. Weirauch, M. T. et al. Determination and inference of eukaryotic transcription factor sequence specificity. Cell 158, 1431–1443 (2014).

89. JASPAR 2022: the 9th release of the open-access database of transcription factor binding profiles | Nucleic Acids Research | Oxford Academic. https://academic.oup.com/nar/article/50/D1/D165/6446529?login=true.

90. Kulkarni, S. R., Jones, D. M. & Vandepoele, K. Enhanced Maps of Transcription Factor Binding Sites Improve Regulatory Networks Learned from Accessible Chromatin Data. Plant Physiol 181, 412–425 (2019).

91. Grant, C. E., Bailey, T. L. & Noble, W. S. FIMO: scanning for occurrences of a given motif. Bioinformatics 27, 1017–1018 (2011).

92. Frith, M. C., Li, M. C. & Weng, Z. Cluster-Buster: finding dense clusters of motifs in DNA sequences. Nucleic Acids Res 31, 3666–3668 (2003).

93. Santana-Garcia, W. et al. RSAT 2022: regulatory sequence analysis tools. Nucleic Acids Research 50, W670–W676 (2022).

94. Bailey, T. L. STREME: accurate and versatile sequence motif discovery. Bioinformatics 37, 2834–2840 (2021).

95. Biswas, A. & Narlikar, L. A universal framework for detecting cis-regulatory diversity in DNA regions. Genome Res 31, 1646–1662 (2021).

96. Smet, D., Opdebeeck, H. & Vandepoele, K. Predicting transcriptional responses to heat and drought stress from genomic features using a machine learning approach in rice. Front. Plant Sci. 14, (2023).

97. Pedregosa, F. et al. Scikit-learn: Machine Learning in Python. Journal of Machine Learning Research 12, 2825–2830 (2011).

98. Akiba, T., Sano, S., Yanase, T., Ohta, T. & Koyama, M. Optuna: A Next-generation Hyperparameter Optimization Framework. in Proceedings of the 25th ACM SIGKDD International Conference on Knowledge Discovery & Data Mining 2623–2631 (Association for Computing Machinery, New York, NY, USA, 2019). doi:10.1145/3292500.3330701.

99. Lundberg, S. M. & Lee, S.-I. A unified approach to interpreting model predictions. in Proceedings of the 31st International Conference on Neural Information Processing Systems 4768–4777 (Curran Associates Inc., Red Hook, NY, USA, 2017).

100. Ferrari, C., Manosalva Pérez, N. & Vandepoele, K. MINI-EX: Integrative inference of single-cell gene regulatory networks in plants. Mol Plant 15, 1807–1824 (2022).

101. Shannon, P. et al. Cytoscape: a software environment for integrated models of biomolecular interaction networks. Genome Res 13, 2498–2504 (2003).

102. Jones, P. et al. InterProScan 5: genome-scale protein function classification. Bioinformatics 30, 1236–1240 (2014).

103. Roberts, W. R., Ruck, E. C., Downey, K. M., Pinseel, E. & Alverson, A. J. Resolving Marine–Freshwater Transitions by Diatoms Through a Fog of Gene Tree Discordance. Systematic Biology syad038 (2023) doi:10.1093/sysbio/syad038.

