## Supplementary Materials for "A Myb-dominated gene regulatory network universally controls sexual cell fate transitions in diatoms"

* Current address: Cell Biology and Biophysics Unit, EMBL, Heidelberg, Germany
### Current address: Laboratory of Cell and Developmental Biology, Wageningen University, Wageningen, The Netherlands
** Current address: Laboratory of Cell Cycles of Algae, Centre Algatech, Institute of Microbiology of the Czech Academy of Sciences, Trebon, Czech Republic

### Supplementary Figures

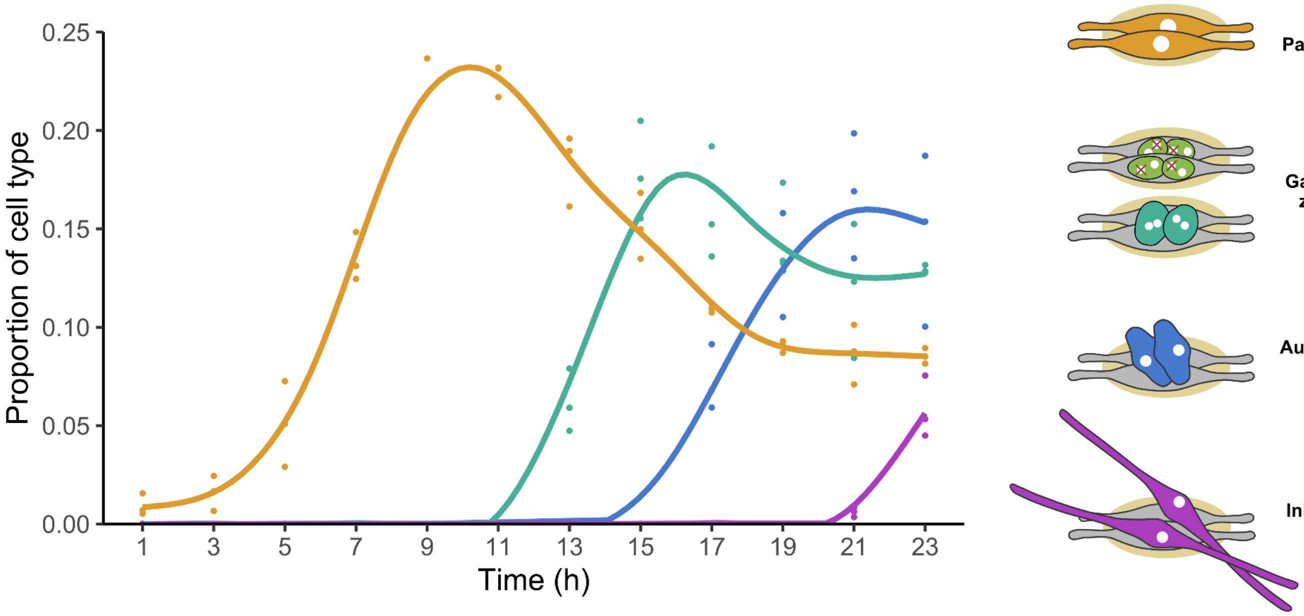


**Figure S1: Reference time series of sexual reproduction in the diatom *Cylindrotheca closterium*.** The proportion of different sexual cell types (colors) is shown in function of the time since compatible *C. closterium* strains ZW2.20 & CA1.15 were crossed. Points reflect individual data points while smooth lines connect averages over time using a generalized additive model (GAM).

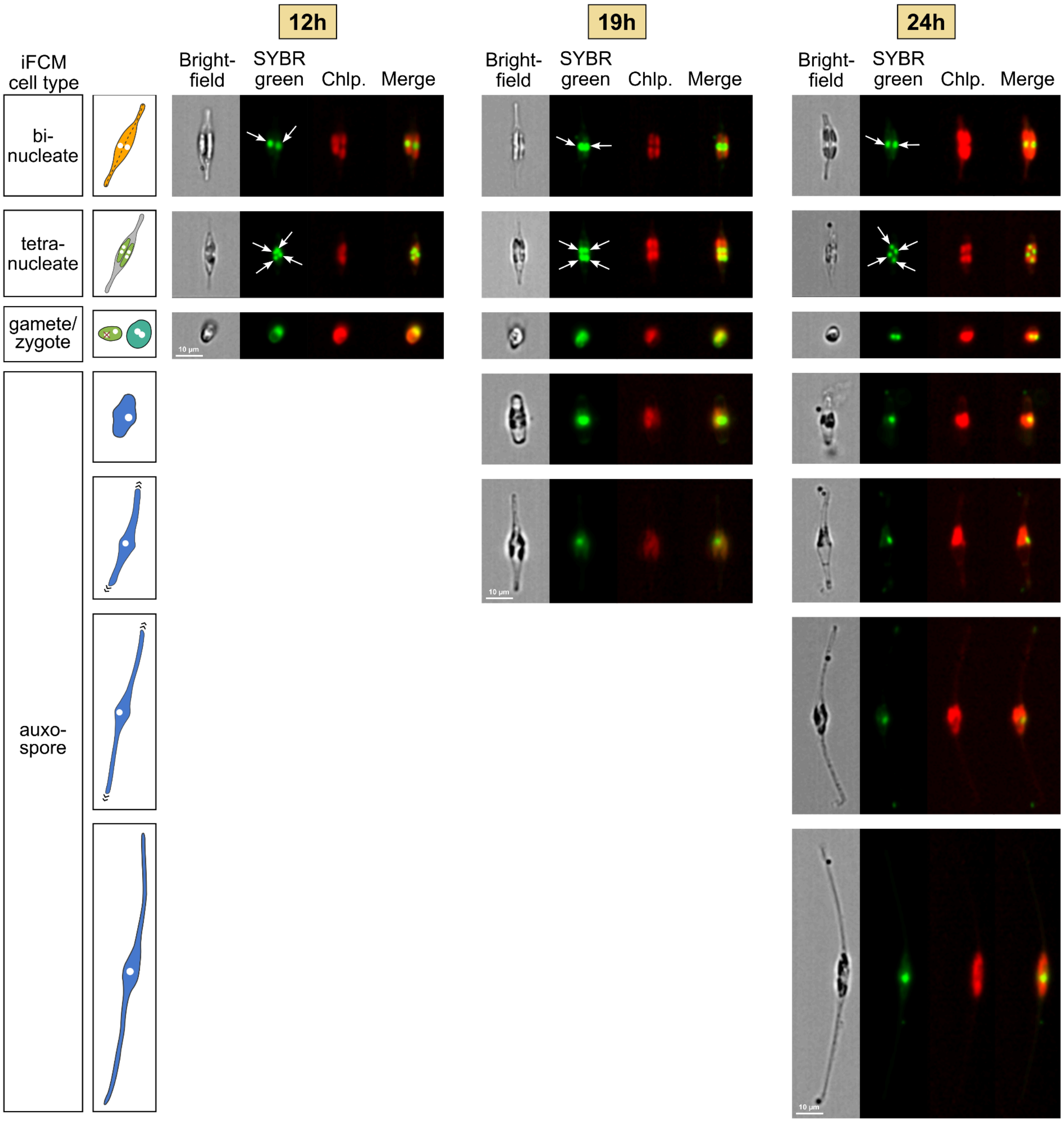


**Figure S2: Imaging flow cytometry of cell cycle progression in different sexual cell types.** Microscopic pictures from three samples (12h, 19h and 24h post-cross) showing different cell types identified using imaging flow cytometry (iFCM) by an Amnis ImageStream X Mark II machine. Images were obtained in brightfield, SYBR green and chloroplast autofluorescence (Chlp.) channels. The latter two were combined in the "Merge" channel. White arrows indicate individual nuclei.

| | 12h | 19h | 24h | Combined |
|---|---|---|---|---|
| # raw PE reads | 118,958,562 | 450,829,637 | 153,488,490 | 723,276,689 |
| # cellular reads | 60,605,029 | 318,623,460 | 72,361,878 | 451,590,367 |
| # high-quality cells | 1079 | 3448 | 4147 | 8674 |
| Total # UMIs | 1,876,062 | 8,477,939 | 7,298,524 | 17,652,525 |
| Avg. # UMIs per cell | 1739 | 2459 | 1760 | 2035 |
| Total # of genes expr. | 18,027 | 20,238 | 19,371 | 20,534 |
| Avg. # genes per cell | 1044 | 1392 | 1073 | 1196 |

**Figure S3:** Table showing scRNA-seq statistics obtained at three time points following the mixture of two mating types (12h, 19h and 24h post-cross), as well as the merged object ("Combined").

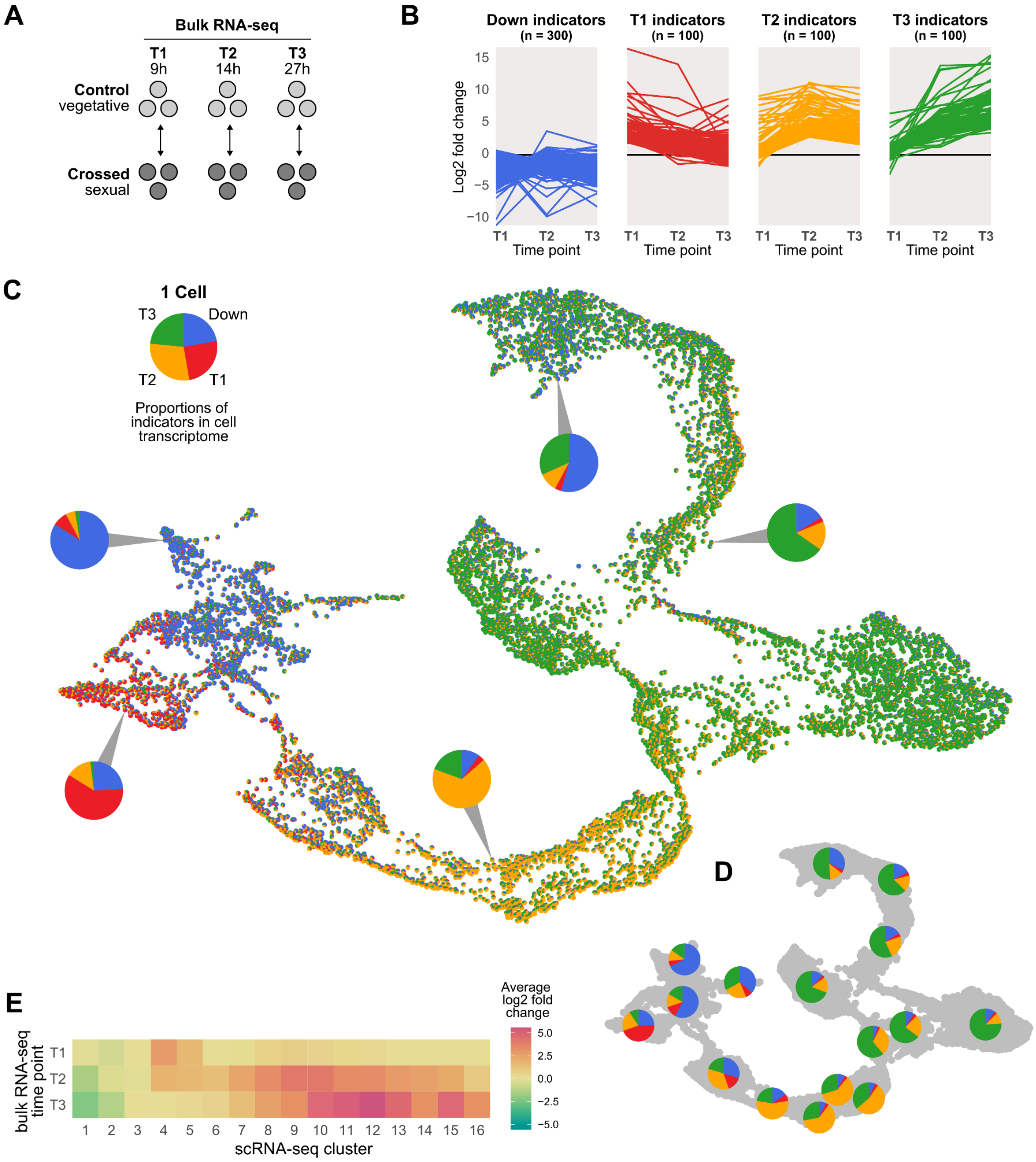


**Figure S4: Annotating single-cell developmental trajectory with bulk RNA-seq time series data.**

**(a)** Experimental set-up of bulk RNA-seq experiment of Audoor et al. 2024. **(b)** Temporal response of four classes of indicator genes in the bulk RNA-seq time series experiment: vegetative (Down), early (T1), middle (T2) and late stages (T3) of sexual reproduction. Line plots show the log2 fold changes in sexual relative to vegetative conditions over time. **(c)** Pie charts showing the relative expression of indicator genes in the transcriptome of a single cell are laid out in a UMAP plot. Example pie charts are enlarged. **(d)** Pie charts showing the relative expression of indicator genes, averaged for each of the 16 single-cell clusters, **(e)** Heatmap of the average differential expression (log2 fold changes) of top-20 clustering-based marker genes for each scRNA-seq cluster in the three bulk RNA-seq time points.

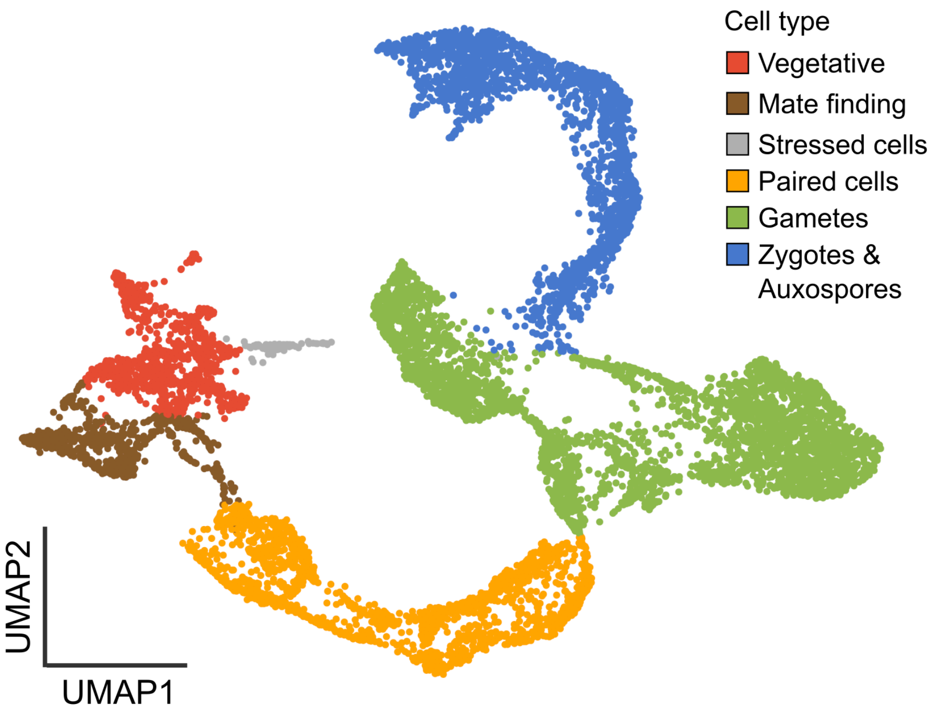


**Figure S5: Consensus cell type identification of 8674 single cells.**
Cell type annotation was based on a combination of cell clustering, expression of gene sets with a known function, correlation with bulk RNA-seq experiments, the temporal information of different scRNA-seq samples and functional validation through reporter lines.

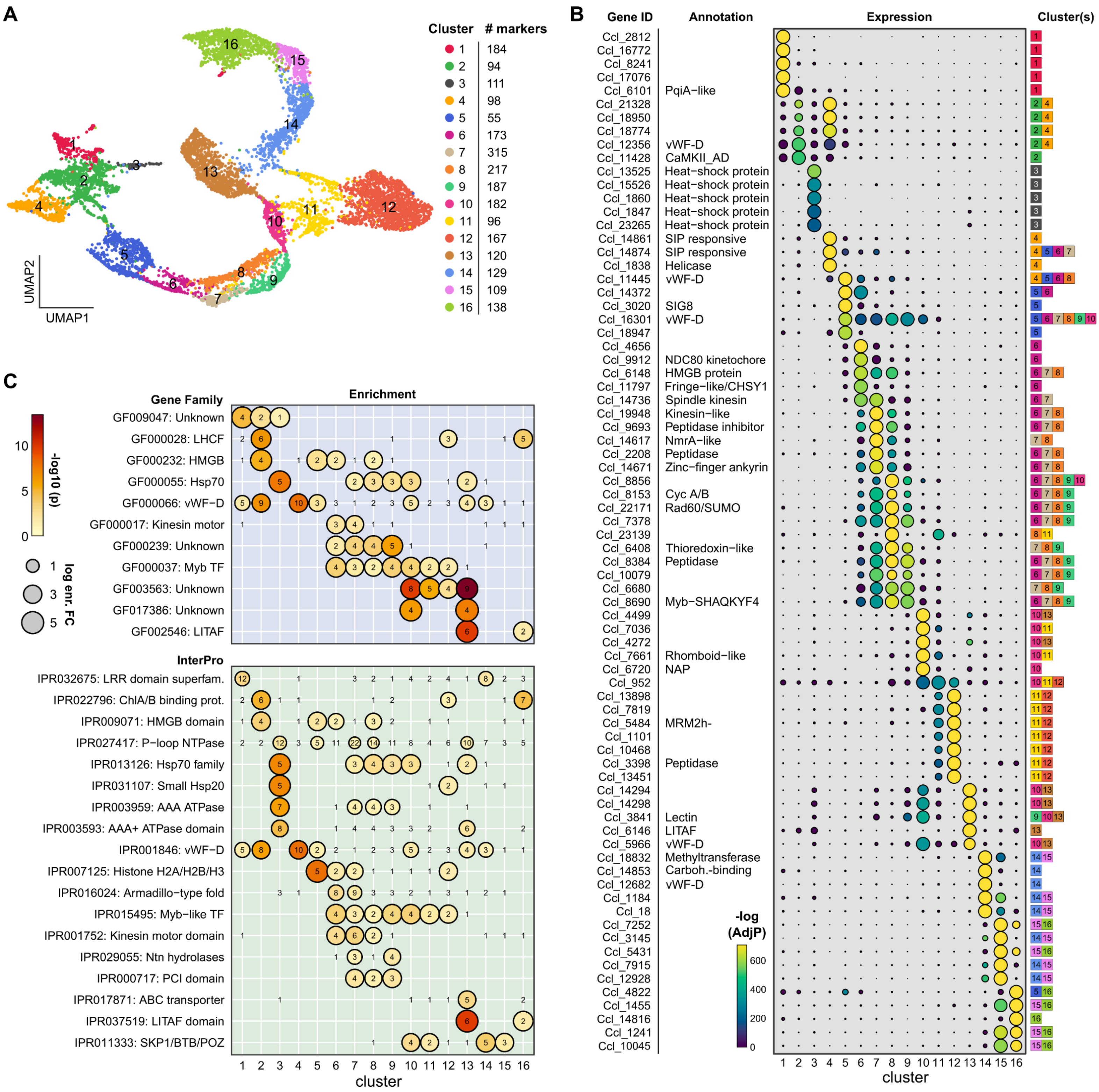

A
Cluster
### markers
1 184
2 94
3 111
4 98
5 55
6 173
7 315
8 217
9 187
10 182
11 96
12 167
13 120
14 129
15 109
16 138
UMAP1
UMAP2
B
Gene ID
Annotation
Expression
Cluster(s)
Ccl_2812
Ccl_16772
Ccl_8241
Ccl_17076
Ccl_6101 PqiA−like
Ccl_21328
Ccl_18950
Ccl_18774
Ccl_12356 vWF−D
Ccl_11428 CaMKII_AD
Ccl_13525 Heat−shock protein
Ccl_15526 Heat−shock protein
Ccl_1860 Heat−shock protein
Ccl_1847 Heat−shock protein
Ccl_23265 Heat−shock protein
Ccl_14861 SIP responsive
Ccl_14874 SIP responsive
Ccl_1838 Helicase
Ccl_11445 vWF−D
Ccl_14372
Ccl_3020 SIG8
Ccl_16301 vWF−D
Ccl_18947
Ccl_4656
Ccl_9912 NDC80 kinetochore
Ccl_6148 HMGB protein
Ccl_11797 Fringe−like/CHSY1
Ccl_14736 Spindle kinesin
Ccl_19948 Kinesin−like
Ccl_9693 Peptidase inhibitor
Ccl_14617 NmrA−like
Ccl_2208 Peptidase
Ccl_14671 Zinc−finger ankyrin
Ccl_8856
Ccl_8153 Cyc A/B
Ccl_22171 Rad60/SUMO
Ccl_7378
Ccl_23139
Ccl_6408 Thioredoxin−like
Ccl_8384 Peptidase
Ccl_10079
Ccl_6680
Ccl_8690 Myb−SHAQKYF4
Ccl_4499
Ccl_7036
Ccl_4272
Ccl_7661 Rhomboid−like
Ccl_6720 NAP
Ccl_952
Ccl_13898
Ccl_7819
Ccl_5484 MRM2h-
Ccl_1101
Ccl_10468
Ccl_3398 Peptidase
Ccl_13451
Ccl_14294
Ccl_14298
Ccl_3841 Lectin
Ccl_6146 LITAF
Ccl_5966 vWF−D
Ccl_18832 Methyltransferase
Ccl_14853 Carboh.-binding
Ccl_12682 vWF−D
Ccl_1184
Ccl_18
Ccl_7252
Ccl_3145
Ccl_5431
Ccl_7915
Ccl_12928
Ccl_4822
Ccl_1455
Ccl_14816
Ccl_1241
Ccl_10045
-log (AdjP)
600
400
200
0
cluster
C
Gene Family
Enrichment
GF009047: Unknown
GF000028: LHCF
GF000232: HMGB
GF000055: Hsp70
GF000066: vWF−D
GF000017: Kinesin motor
GF000239: Unknown
GF000037: Myb TF
GF003563: Unknown
GF017386: Unknown
GF002546: LITAF
-log10 (p)
10
5
0
log enr. FC
1
3
5
InterPro
IPR032675: LRR domain superfam.
IPR022796: ChlA/B binding prot.
IPR009071: HMGB domain
IPR027417: P−loop NTPase
IPR013126: Hsp70 family
IPR031107: Small Hsp20
IPR003959: AAA ATPase
IPR003593: AAA+ ATPase domain
IPR001846: vWF−D
IPR007125: Histone H2A/H2B/H3
IPR016024: Armadillo−type fold
IPR015495: Myb−like TF
IPR001752: Kinesin motor domain
IPR029055: Ntn hydrolases
IPR000717: PCI domain
IPR017871: ABC transporter
IPR037519: LITAF domain
IPR011333: SKP1/BTB/POZ
cluster

**Figure S6: Clustering, identification and functional enrichment of marker genes.**
**(a)** Uniform Manifold Approximation and Projection (UMAP) plot of 8674 *C. closterium* cells in the combined dataset. Points are coloured according to the cell cluster they belong to. The number of marker genes identified for each cluster is shown in the legend. **(b)** Dotplot showing the expression of the top 5 most significant marker genes for each cluster. The point size shows the average expression of a gene in a cluster, normalized between one (maximal expression) and zero (no expression). The -log adjusted p-value of marker differential expression in each cluster is shown by the dots' color. Genes that were not markers for a given cluster have a p-value of 1. **(c)** Dotplots showing enrichment of homologous gene families and InterPro domains as features (y-axis) among the marker genes of each cluster (x-axis). Points vary in size with the enrichment fold change (FC) and are coloured according to the p-value of enrichment (hypergeometric test). Dots are only shown for clusters in which a feature is significantly enriched. The number of genes that are among the marker genes of a cluster are plotted on top.

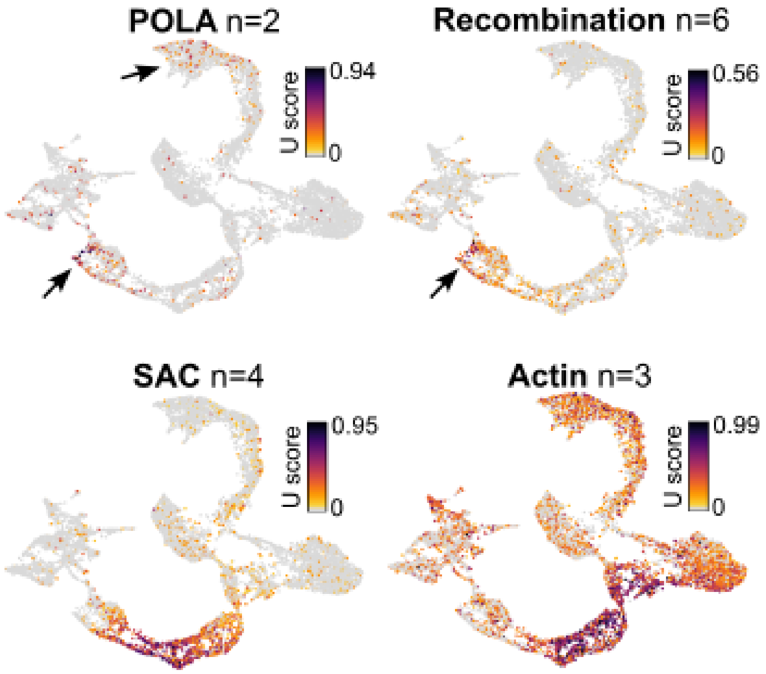


**Figure S7: Expression of cell cycle markers.** UMAP plots showing the Mann-Whitney U statistic of cell cycle marker gene sets. Black arrows highlight locations with expression that are hard to distinguish. POLA: DNA polymerase alpha subunits A1 and A2. SAC: spindle assembly checkpoint. “n” shows the number of genes in each set.

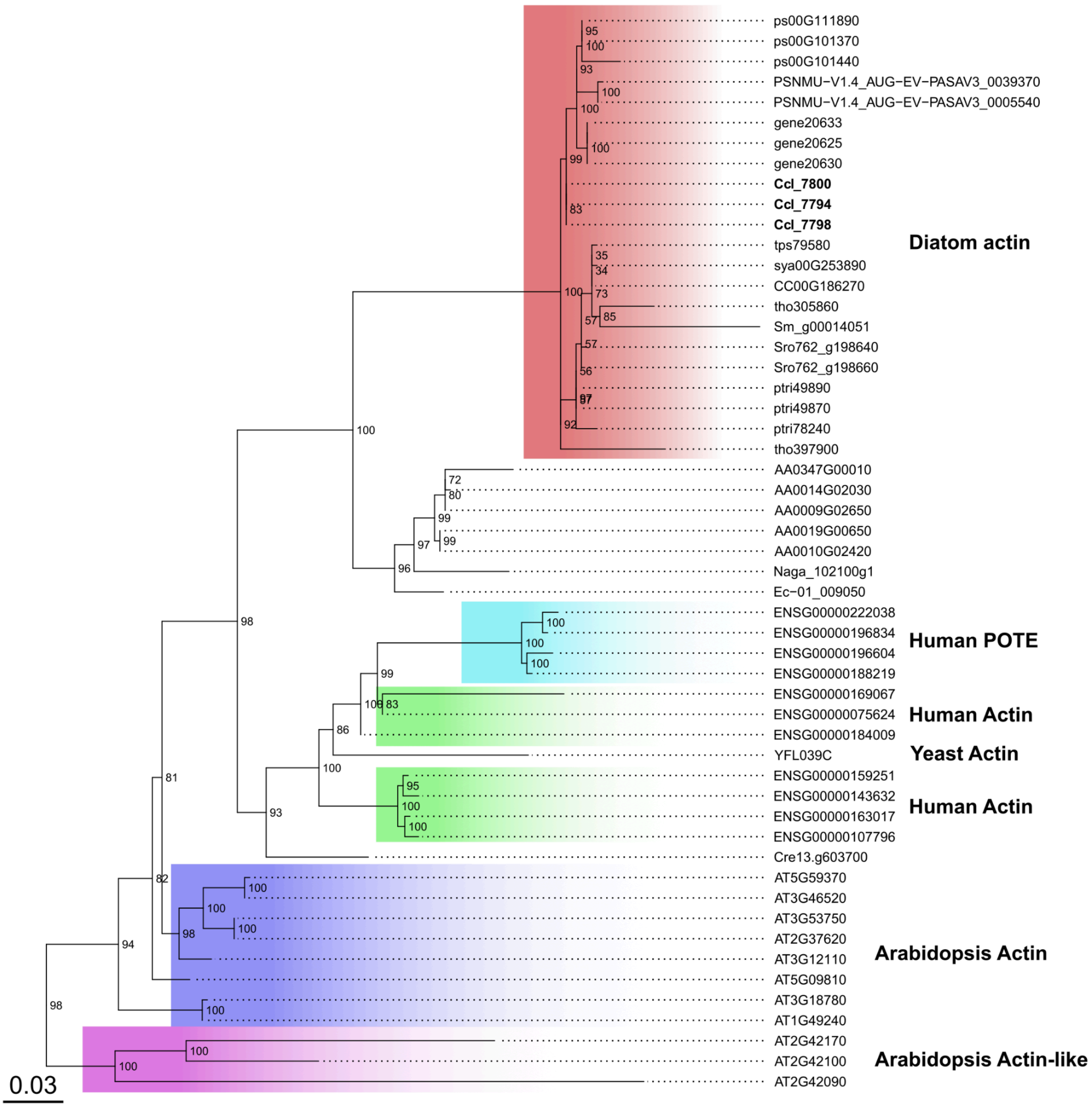


**Figure S8: Bootstrap-consensus phylogenetic tree of the Actin protein family in eukaryotes.** Node labels represent bootstrap support and the tree was midpoint rooted. *C. closterium* actins are indicated in bold.

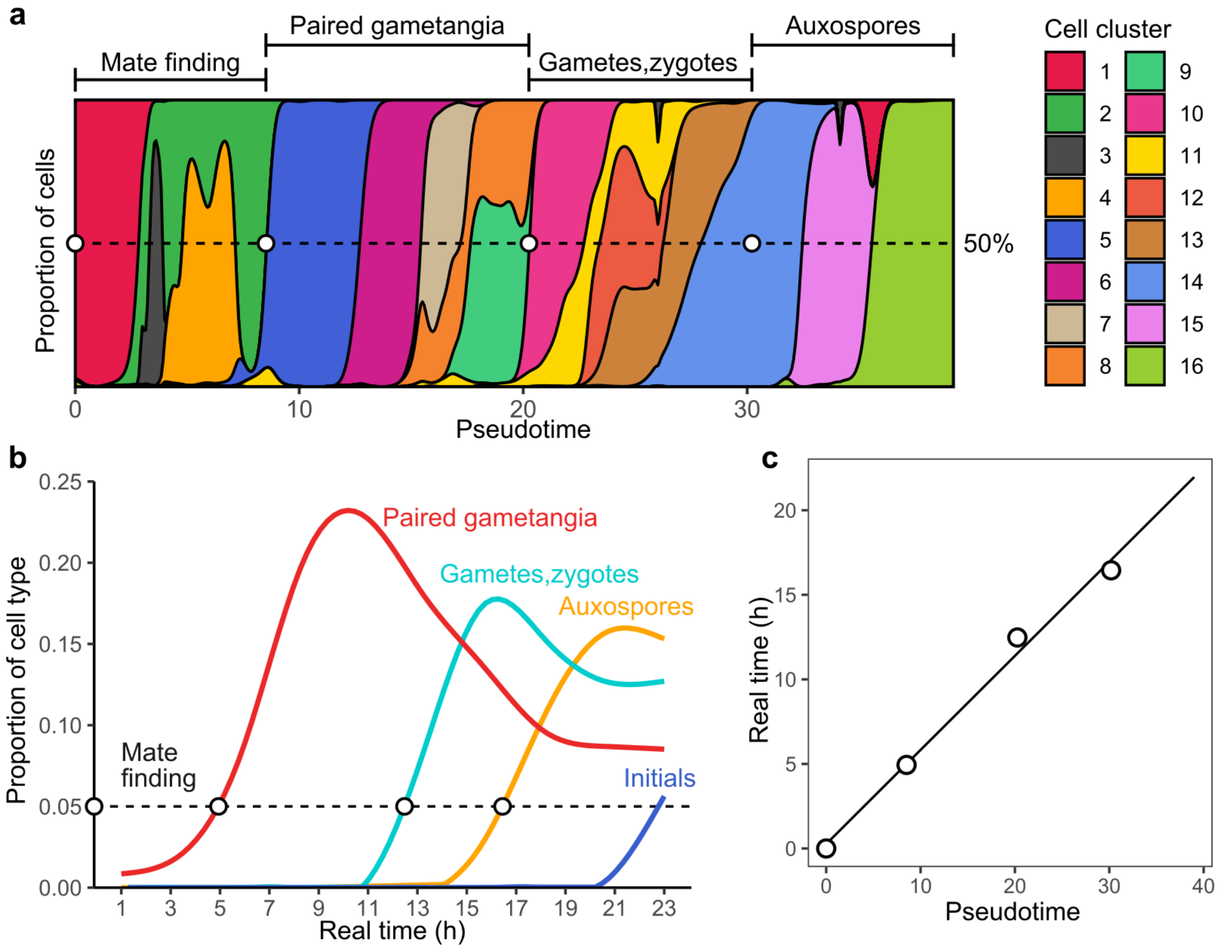


**Figure S9: Correlating pseudotime with real developmental time. (a)** Density plot showing the proportion of cells in the scRNA-seq atlas belonging to each cluster in function of pseudotime. Pseudotime intervals are shown on the top, bordered by empty circles. **(b)** Smoothed curve of cell types appearing after a synchronized cross. Time intervals between the appearance of 5% of each new cell type are bordered by empty circles. **(c)** Dot plot comparing the appearance of new cell types in pseudotime (scRNA-seq) with real time (laboratory cross). The line represents the linear model fit on the data.

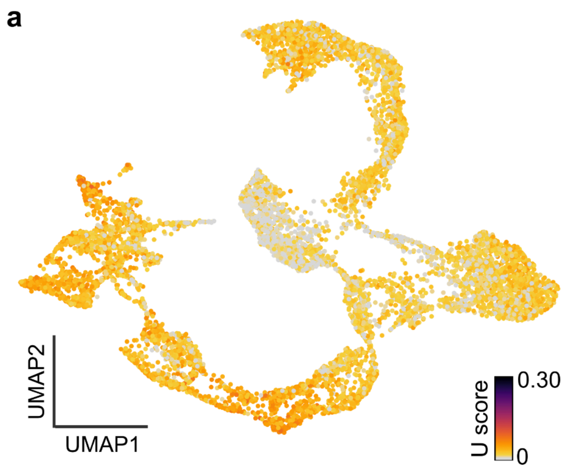


**Figure S11: UMAP plot showing the Mann-Whitney U statistic of each cell for bulk RNA-seq gene sets.**

(**a**) Upregulated genes after 3h of sex-inducing pheromone minus (SIP-) treatment (n = 207), (**b**) Upregulated genes after 9h of SIP- treatment (n = 65).

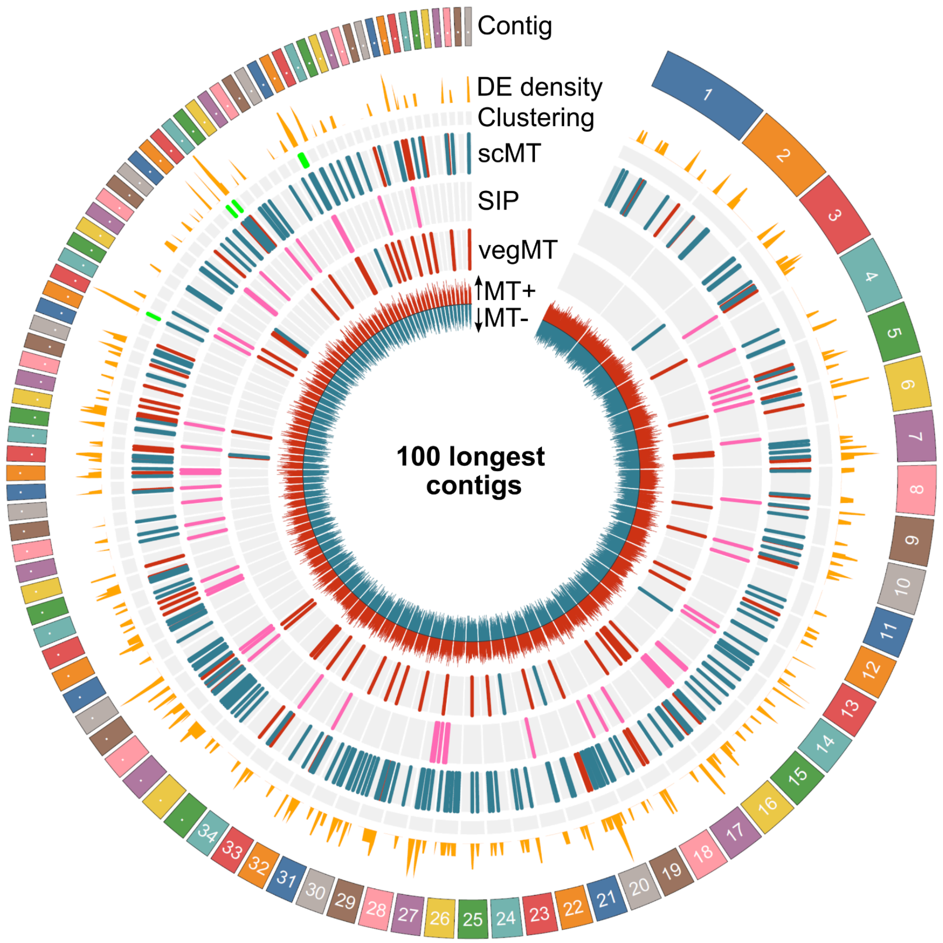


**Figure S12: Circos plot showing functional clustering (in green) of mating type-biased genes along the 100 longest contigs in the *C. closterium* genome.**

Three types of differential expression are shown as concentric bands: between mating types in scRNA-seq data (scMT), in response to sex inducing pheromone (SIP) and between mating types in vegetative cells of six different genotypes (vegMT).

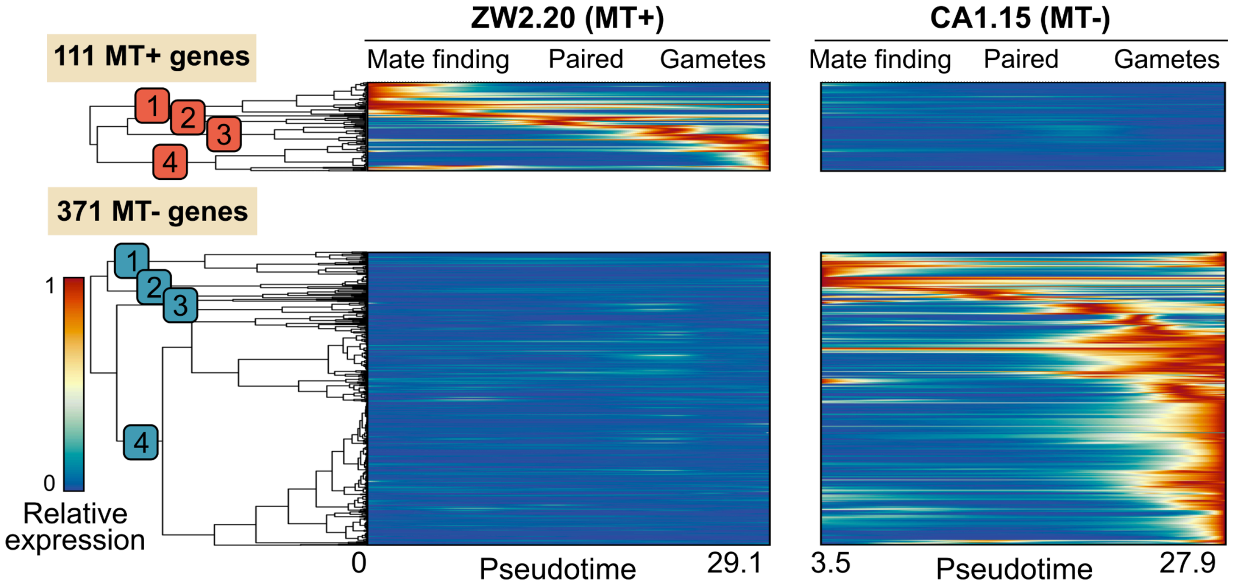


**Figure S13: Heatmaps showing the relative expression of significant MT+ (ZW2.20) and MT- (CA1.15) biased genes (y-axis) over pseudotime, smoothed by generalized additive models (GAMs).**

Expression of the same gene along MT+ and MT- trajectories is shown side-by-side, while genes significantly upregulated in MT+ and in MT- are separated top-to-bottom. Hierarchical dendrograms on the left cluster genes into four numbered modules per mating type.

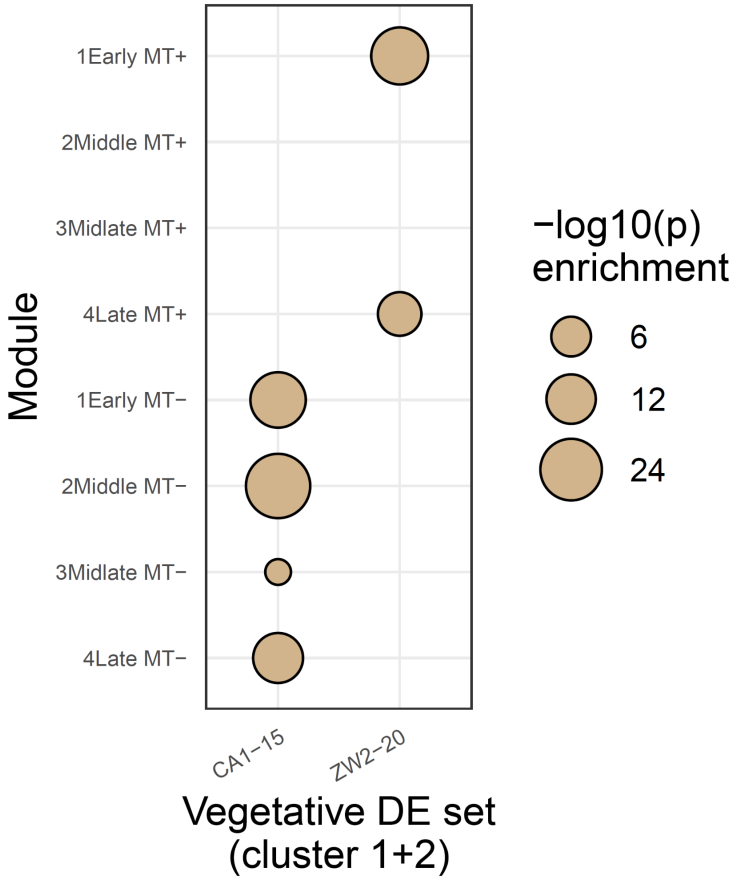


**Figure S14: Bubble plot showing enrichment of genes that are already differentially expressed (DE) in vegetative clusters 1 and 2 in genotype-specific modules.**

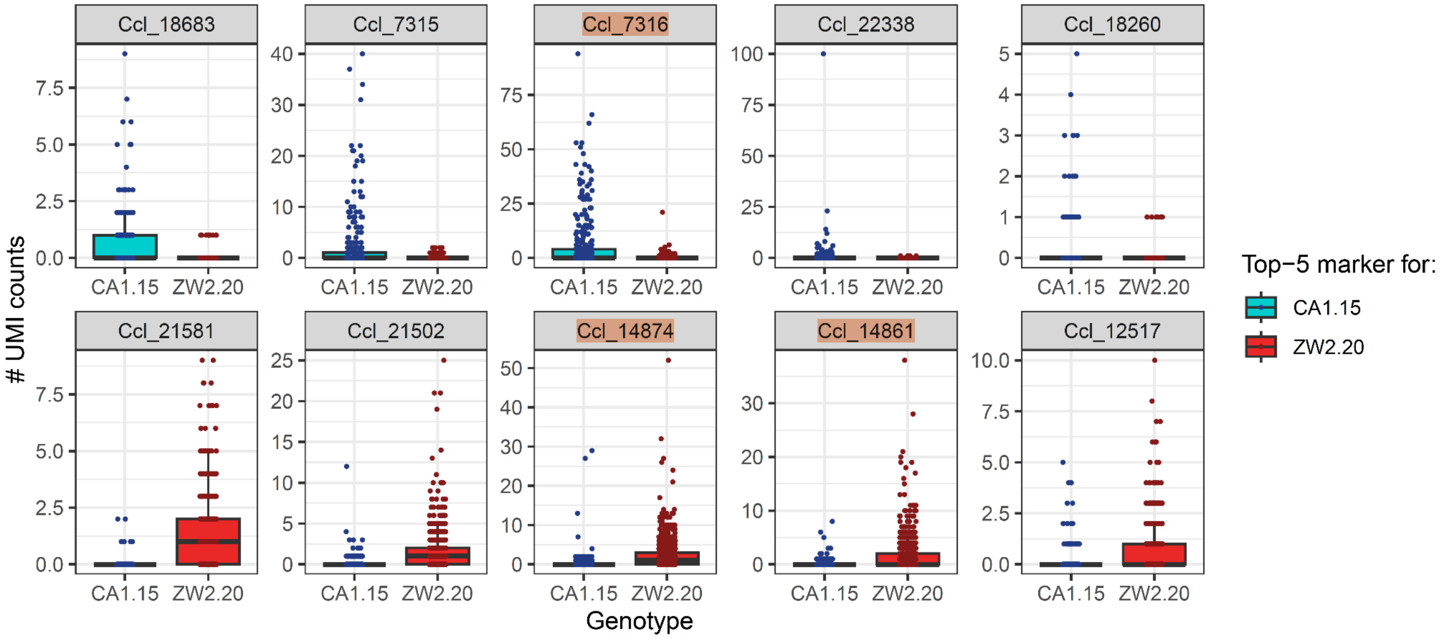


**Figure S15: Expression of top-5 most significant differentially expressed genes between two genotypes (CA1.15 and ZW2.20) in vegetative clusters 1+2, out of a total of 307 differentially expressed genes.**

Each dot represents the expression of a single cell belonging to either genotype (x-axis), summarized as a boxplot. UMI: unique molecular identifier. Vegetative mating type marker genes previously identified by Belisova et al. (2025) are highlighted.

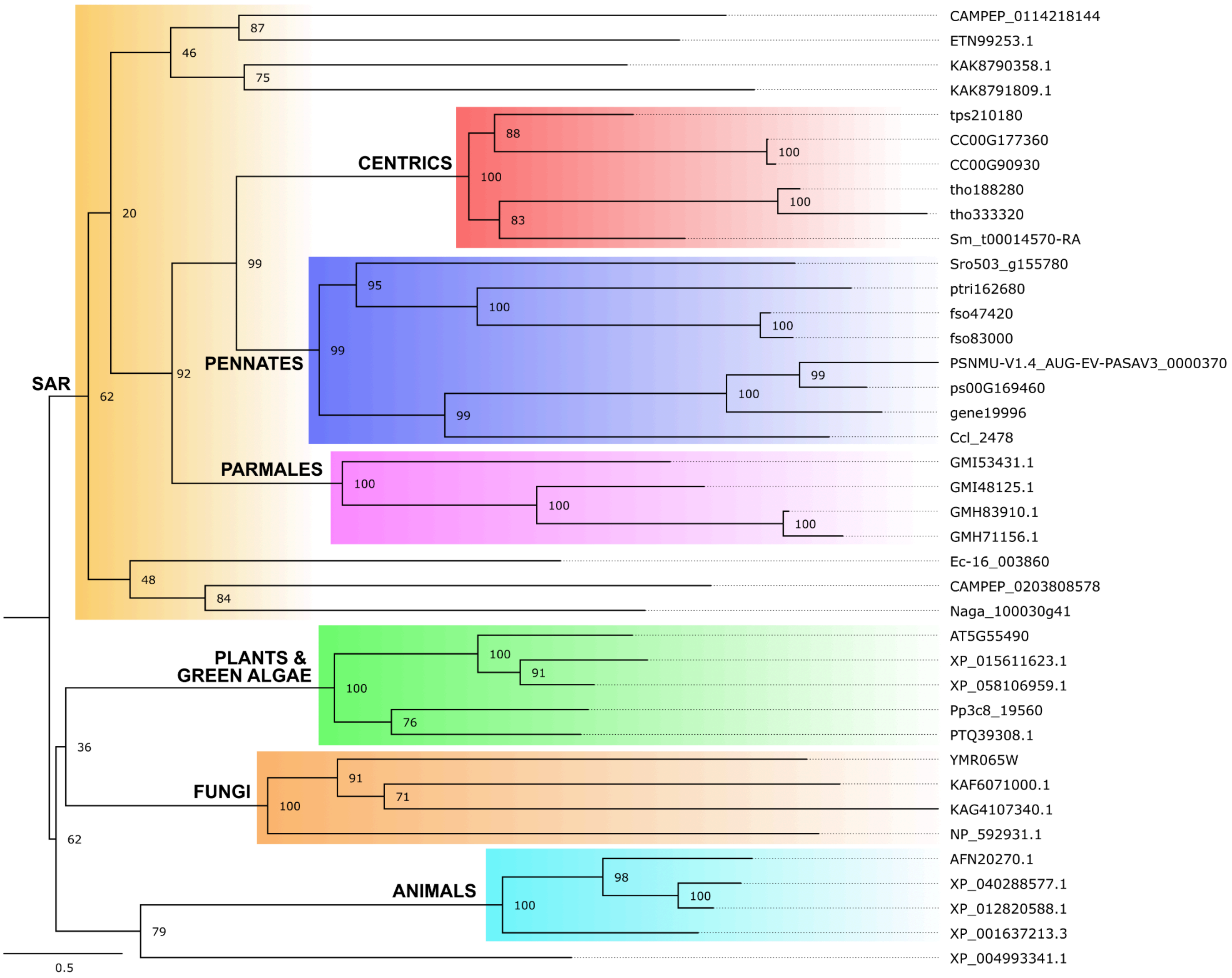


**Figure S16: Midpoint-rooted bootstrap consensus phylogenetic tree of Gamete Expressed Protein 1 (GEX1) proteins in eukaryotes.**

Node labels represent bootstrap support. Protein IDs refer to NCBI, PLAZA Diatoms and MMETSP identifiers.

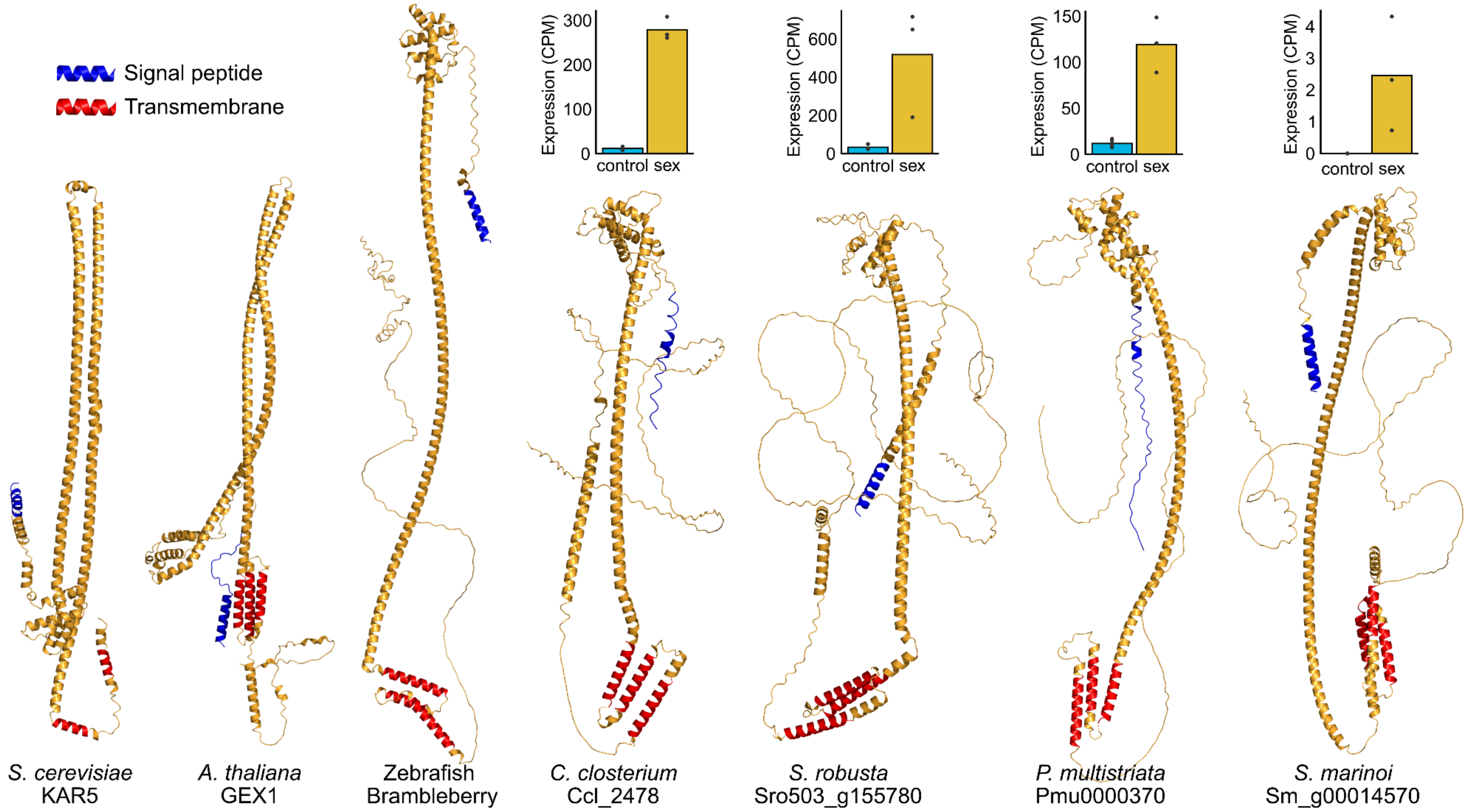


**Figure S17: Alphafold 3 protein structure prediction and bulk transcriptomic expression of GEX1 homologs.**

Bar plots showing expression in counts per million (CPM) of GEX1 homologs during vegetative growth (blue) and sexual reproduction (yellow) in four different diatom species: *Cylindrotheca closterium, Pseudo-nitzschia multistriata, Seminavis robusta* and *Skeletonema marinoi*. Bars show the average expression, while dots indicate individual replicates. CPM: counts per million.

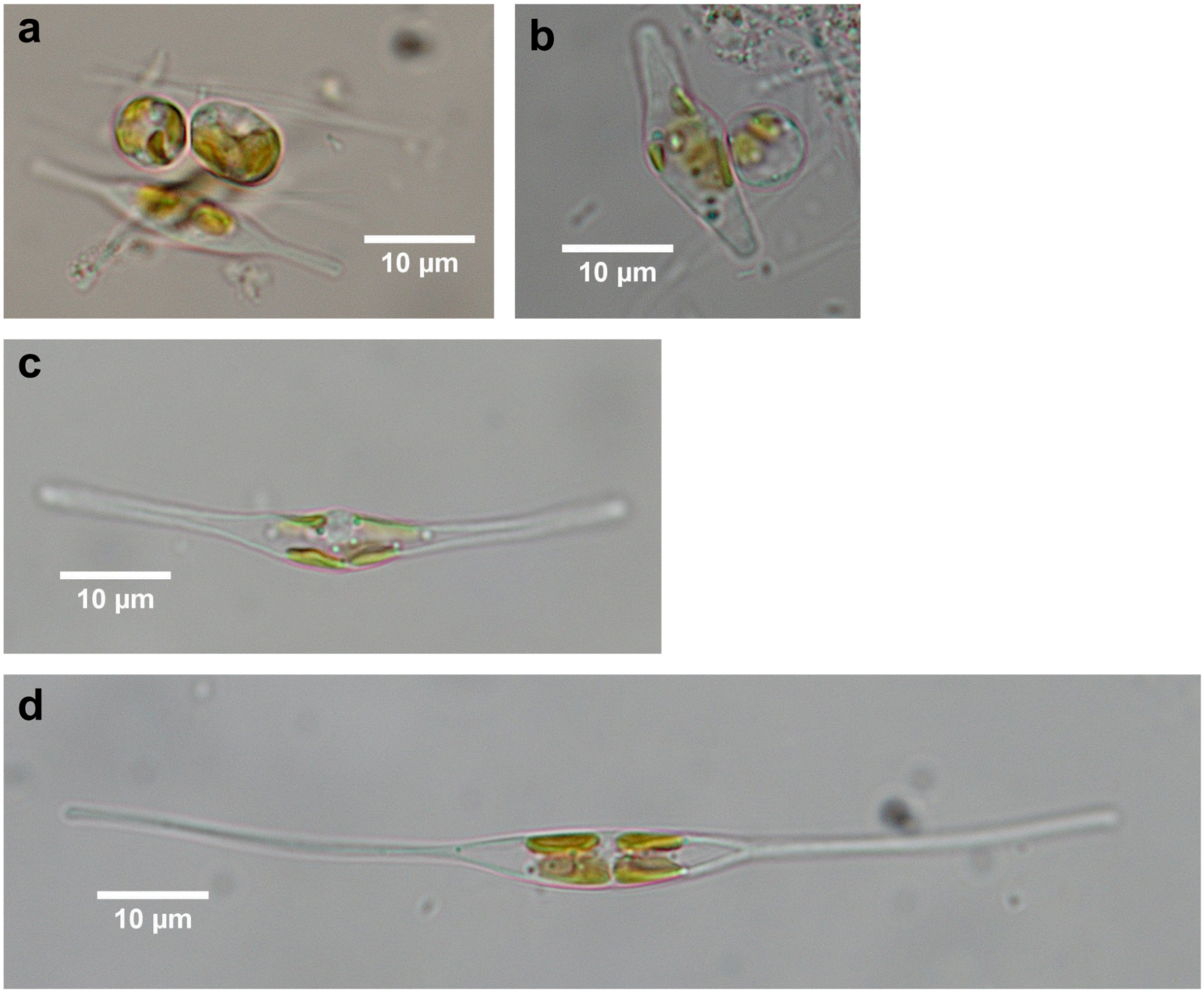


**Figure S18: Auxospore expansion of *C. closterium*, light microscopy.**
**(a)** Two zygotes (round: 6.4 µm diameter, oval: 9.2 µm along longest axis) alongside a vegetative cell, **(b)** A round zygote (7.7 µm diameter) and an expanding young auxospore (22.7 µm length), **(c)** A single expanding auxospore (54.0 µm length) and **(d)** A mature auxospore or initial cell (96.7 µm length).

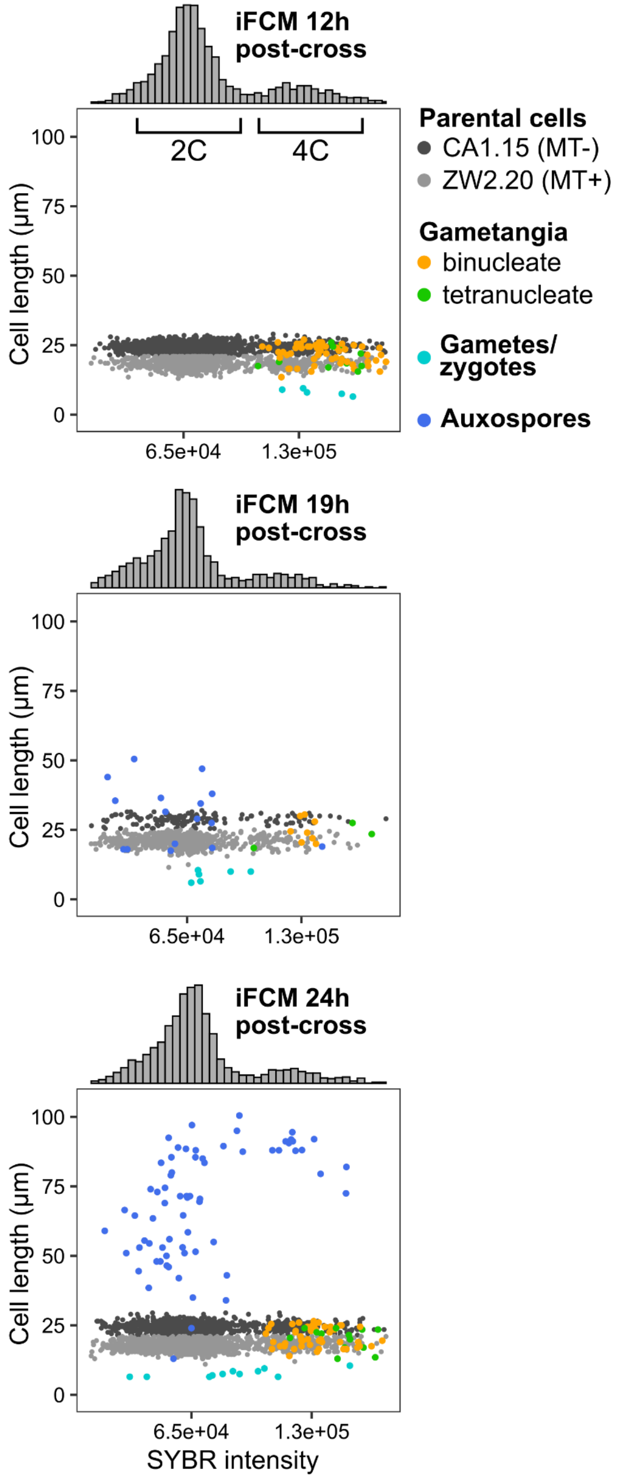


**Figure S19: Scatterplots showing SYBR intensity versus the total cell length, obtained by imaging flow cytometry (iFCM) from a subculture of all three scRNA-seq samples.** Individual cells are coloured according to a microscopy-based cell type identification (nuclei, shape, ploidy, size). The histograms on top summarize the distribution of SYBR intensity of all cells.

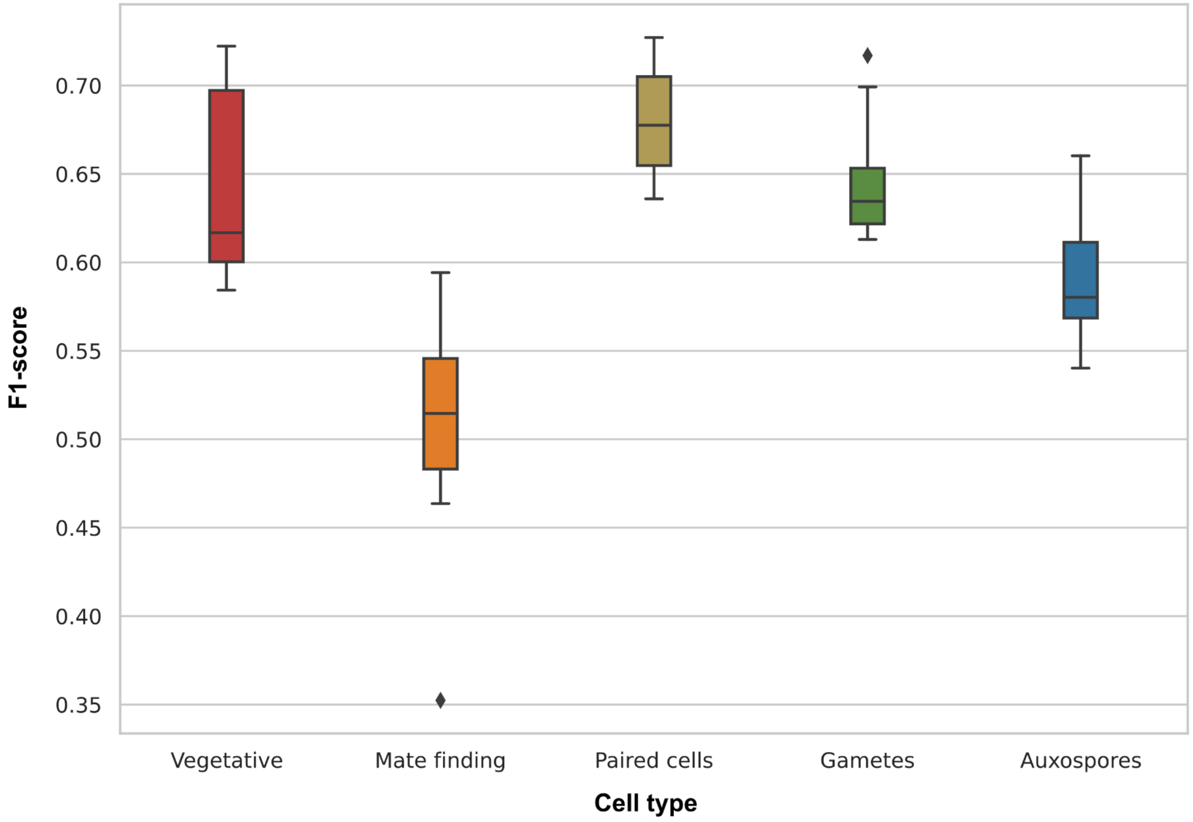


**Figure S20: Performance of Random Forest (RF) classifiers predicting marker genes for five *C. closterium* cell stages based on promoter motif content.**

Motif occurrences in 500-bp upstream regions were used as features in RF models. Each boxplot shows F1-scores from 10 replicate models trained with different train-test splits in function of cell type: vegetative, mate-finding, paired cells, gametes, and auxospores.

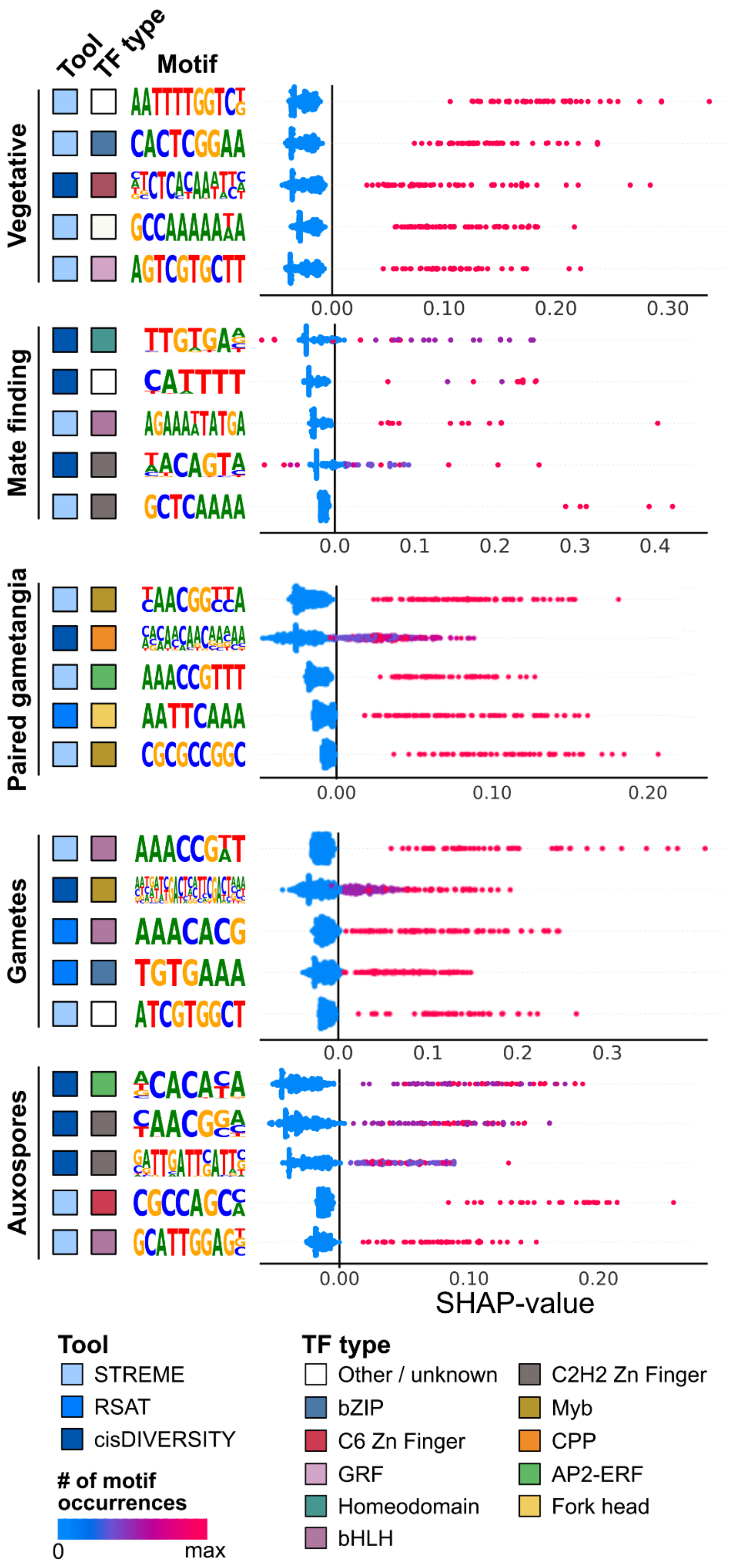
Tool
TF type
Motif
Vegetative
Mate finding
Paired gametangia
Gametes
Auxospores
0.00
0.10
0.20
0.30
0.0
0.1
0.2
0.3
0.4
0.00
0.10
0.20
0.0
0.1
0.2
0.3
0.00
0.10
0.20
SHAP-value
Tool
STREME
RSAT
cisDIVERSITY
### of motif occurrences
0
max
TF type
Other / unknown
bZIP
C6 Zn Finger
GRF
Homeodomain
bHLH
C2H2 Zn Finger
Myb
CPP
AP2-ERF
Fork head

**Figure S21: Key promoter motifs predictive of marker gene identity across five *C. closterium* cell types, determined by SHAP analysis of Random Forest models.**
SHAP (SHapley Additive exPlanations) analysis summary plots displaying the impact (x-axis) of each motif (y-axis) on individual promoter predictions (dots), with colors indicating the number of motif occurrences per promoter. Positive SHAP values indicate a contribution toward the positive class prediction (i.e., marker gene identity). For each cell type, the top-5 ranked motifs are shown, annotated by the tool of discovery in the 500-bp upstream promoter region ("Tool") and the predicted transcription factor type by TomTom ("TF type").

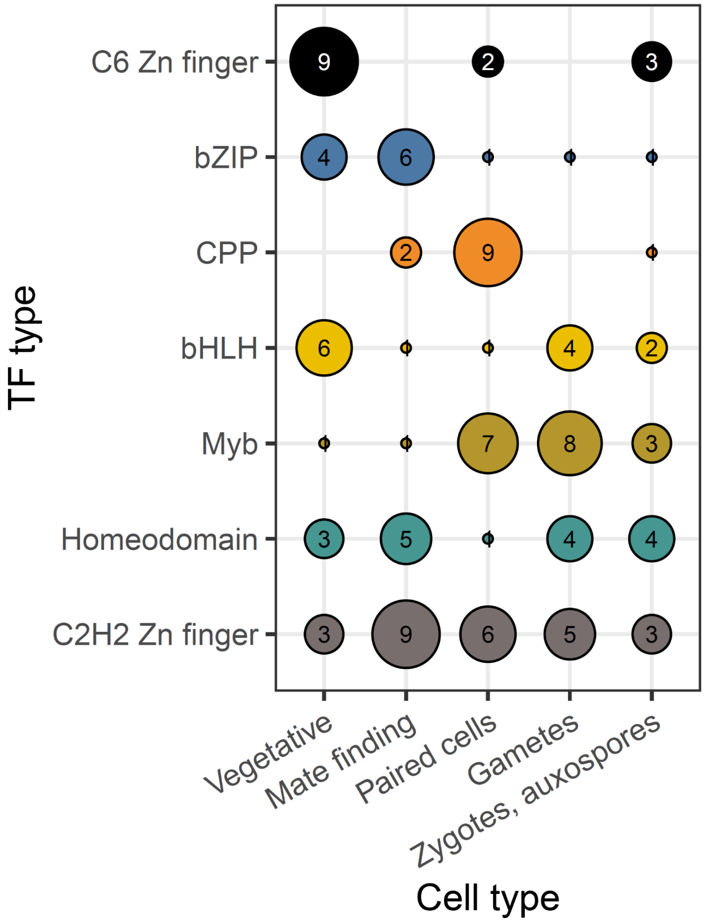


**Figure S22: Robustness of machine learning models in predicting motifs associated with transcription factor types.**

The radius of the circles shows the frequency of occurrence of the seven most common transcription factor (TF) types (y-axis) in function of cell type (x-axis) during 10 independent machine learning runs.

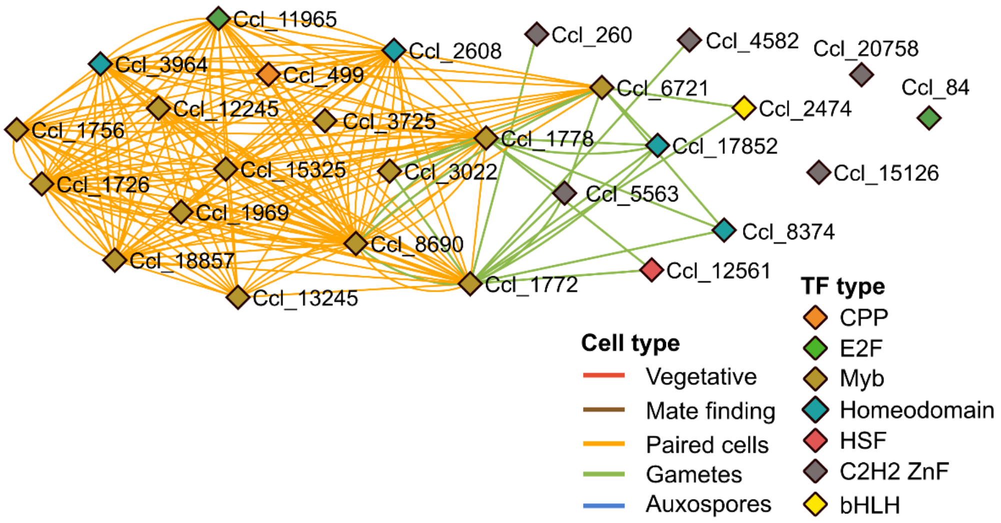


**Figure S23: Subnetwork of gene regulatory network that only shows TF-TF interactions.**
TF: transcription factor. Edges are coloured by the cell type where the regulon was active, diamonds are coloured by the TF type.

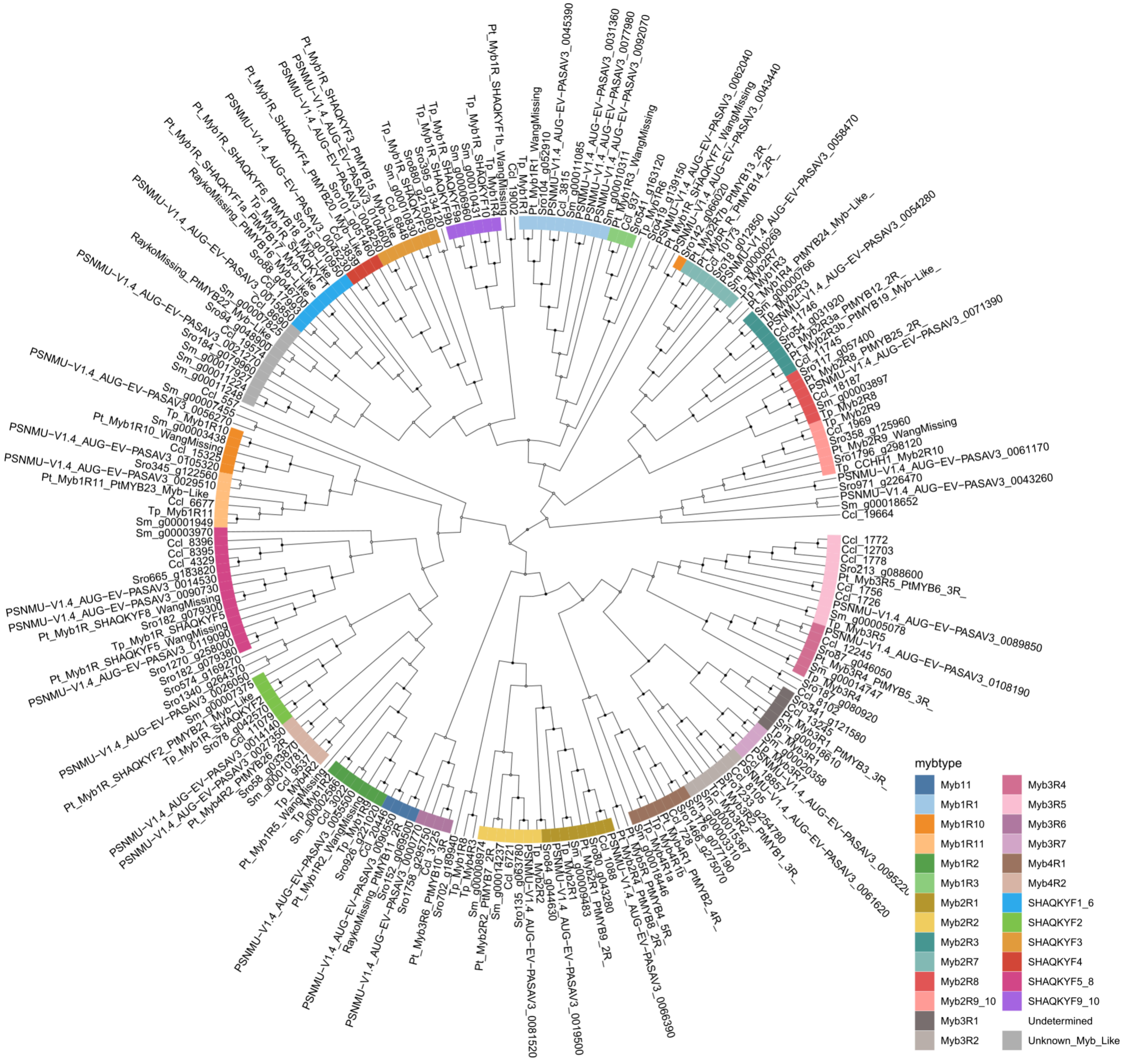


**Figure S24: Midpoint-rooted bootstrap-consensus phylogenetic tree of the Myb transcription factor family in diatoms.** The coloured band indicates the different annotated Myb subtypes.

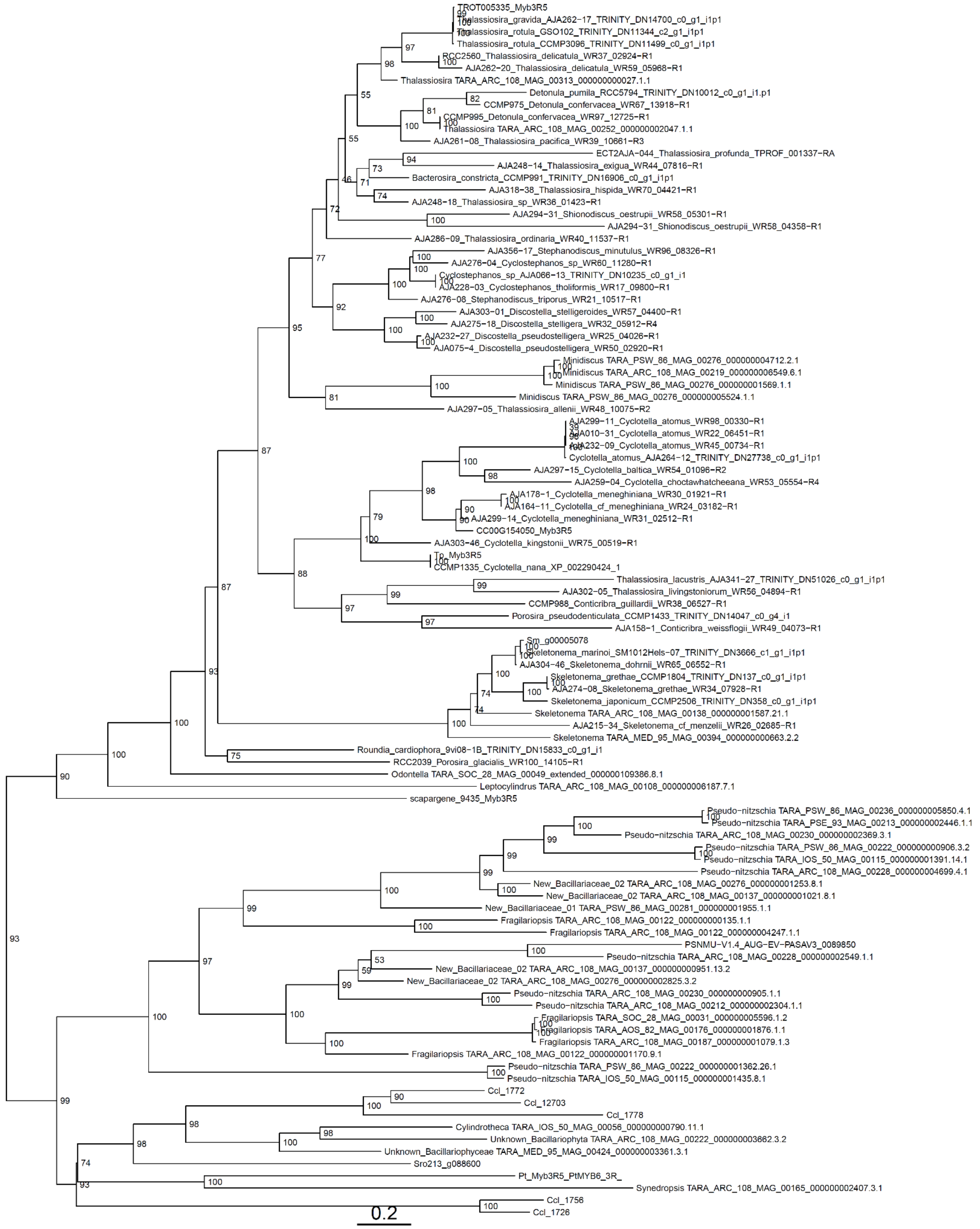


**Figure S25: Midpoint-rooted bootstrap-consensus phylogenetic tree of the Myb3R5 clade hits from Tara Oceans**. This tree includes 36 Myb3R5 hits from Tara Oceans alongside reference Myb3R5 proteins from PLAZA Diatoms, *Cylindrotheca closterium* and Roberts et al. 2023 [1]. Node labels indicate bootstrap support.

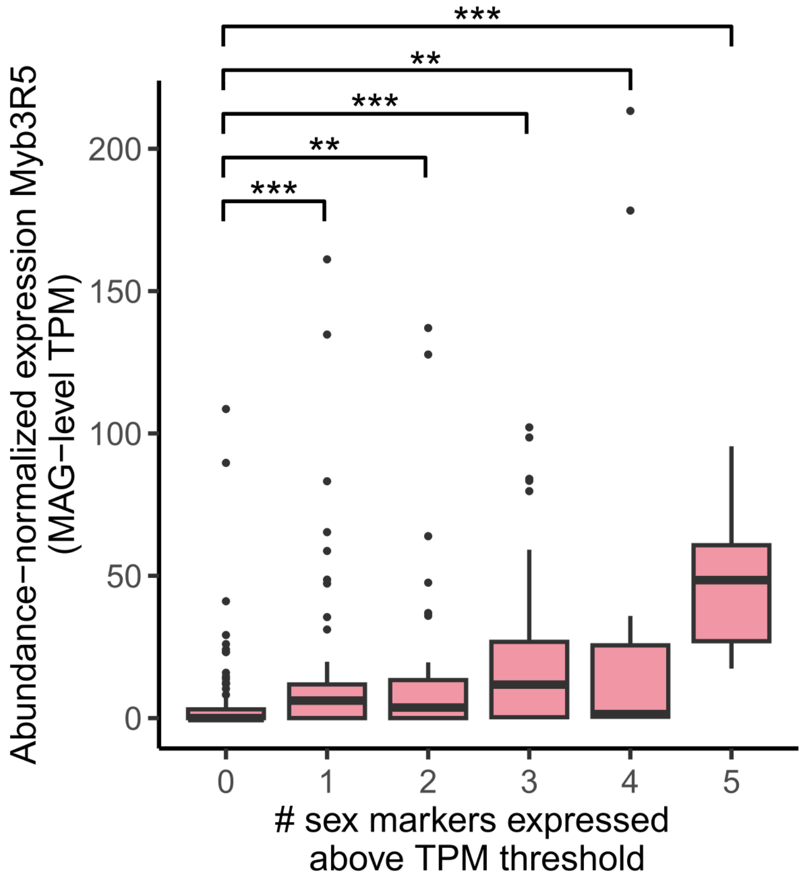


**Figure S26: Boxplots showing the expression of Myb3R5 homologs (in transcripts per million, TPM) in function of the number of sex markers that is expressed relative to a threshold [2].** The central line of the boxplot indicates the median, the box limits show the 25th and 75th percentiles and whiskers extend up to 1.5× the interquartile range. Dots indicate outliers. ** $p < 0.01$, ***: $p < 0.001$

### Supplementary Text

**Text S1: Development of BD Rhapsody-based scRNA-seq for micro-algae**

Although protists have the obvious advantage that no cell dissociation is needed, high-throughput and high-yield scRNA-seq datasets are lacking for most protist taxa due to (1) their specialized cell walls preventing cell lysis, (2) many protists having a small cell volume containing few mRNA molecules, and (3) the difficulty to adapt existing platforms, especially microfluidics, to protist cells with a unique morphology[3–5]. Here, we developed a microwell-based method for diatoms using the BD Rhapsody system, which has several advantages to study protist biology: (1) no protoplastic or nuclear extraction is needed, avoiding stress-related gene expression, (2) cultures can be loaded in their native medium, so there is no osmotic shock, (3) as long as enough sedimentation time is provided - and possibly aided by gentle centrifugation of the cartridge, cell capture should be independent of cell shape, (4) sedimentation and cell density can be verified by automated imaging of the whole cartridge, (5) the Rhapsody protocol is very flexible. Longer lysis times, specialized lysis buffers, chemical removal of the cell wall and freeze-thaw cycles are possible to study recalcitrant protist species.

Our method should, however, be further finetuned for future research. For example, the top marker genes for cluster 3 are heat-shock proteins, suggesting that these cells were stressed during the experiment or culturing, although it is unclear how. Consequently, they were excluded from the analysis of sexual development. Furthermore, our scRNA-seq libraries were markedly depleted in vegetative cells compared to the original cultures, suggesting that sexual cell types lyse more easily, and further optimization might be necessary before high-throughput scRNA-seq of non-sexual diatom cells is feasible.

**Text S2: Sequencing depth and sample size for BD Rhapsody-based scRNA-seq**

To explore the saturation of the library with increasing sequencing depth, we performed additional deep sequencing of the 19h sample. Despite the three-fold higher sequencing depth compared to the other two samples, the average number of UMIs hardly increased, suggesting that 150 million reads is sufficient to capture most of the transcriptome (**Fig. 1d**). In addition, the number of cells was varied to investigate the capture rate and the occurrence of multiplets in microwells. In total, 53, 60 and 98 thousand cells were loaded into the Rhapsody cartridge for 12h, 19h and 24h respectively. After sequencing, mapping and quality filtering, about 4% of these cells were found in the final library. Loading a higher number of cells resulted in a marked increase in the detected number of cells: despite a fivefold lower sequencing depth, we observed a higher number of cells after 24h (4147) compared to 19h (3448). Even when loading almost 100,000 cells, most microwells contained only a single cell after sedimentation. We did observe that cells of the same cell type (paired cells or gametes) can remain attached to each other during sedimentation into the microwells, which resulted in a layer of cells with a recombined genotype in between MT+ and MT- (**Fig. 4a**). Interestingly, the lowest number of cells was detected for the 12h culture, which consisted predominantly of unpaired cells that were still protected by the silica cell wall. Hence, cell lysis in the Rhapsody cartridge may be impeded in more silicified cell types such as vegetative cells.

**Text S3: Determining cell type identity of 16 unknown clusters**

Since there are no reference single-cell RNA-seq datasets of diatom life cycles (or any protist for that matter), we annotated the 16 scRNA-seq clusters using a combination of different methods:

(1) Repeated scRNA-seq sampling at different time intervals. Cells from the 12h-sample were limited to the initial stages during which vegetative cells localize a mate and form mating pairs (*Cluster 1-10*). Meanwhile, the later samples (19h and 24h) comprised the full set of sexual cell types and therefore cover the entire developmental pathway (*Cluster 1-16*).
(2) Integration of bulk RNA-seq time series of sex. The sequence of differentiation can be further deduced by determining for each single-cell the relative expression of genes specific for bulk RNA-seq different time points (**Fig. S4**). Cells at the start of the developmental trajectory mainly express genes that are downregulated during sex (*Cluster 1-2*). These are parental cells that have not committed to sex, as well as naive cells that were added as a spike-in to the 12h sample. As single-cells move along the developmental trajectory, their transcriptome sequentially resembles the bulk RNA-seq samples containing mostly mate finding and paired cells (*Cluster 4-5*), paired cells (*Clusters 6-9*), gametes, zygotes and auxospores (*Clusters 10-15*) and eventually regain their vegetative expression pattern (*Cluster 16*).
(3) Pheromone-response experiment. A bulk RNA-seq experiment treating MT+ cells with Sex-Inducing-Pheromone from MT- (SIP-) revealed that *Cluster 4* represents SIP-conditioned MT+ cells.
(4) Timing of known genes. Based on the timing of putative functionally understood genes for mating type differentiation, karyogamy and auxospore identity (**Fig. S6**), we identified the major cell stages of pair formation (*Cluster 5-9*), gametogenesis/gametes (*Cluster 10-13*) and auxospore expansion (*Cluster 14-16*).
(5) Cell cycle phasing analysis. Expression-based classification of each cell in the scRNA-seq dataset by their cell cycle phase (G1, S, G2/M) showed a clear cell cycle progression in Clusters 5-9, further supporting their annotation as gametangia (**Fig. 2c**). While Cluster 1-4 mostly resided in the G1-phase, cells started expressing DNA replication and recombination programs in Cluster 5, followed by meiotic spindle and chromosome condensation genes in Cluster 6. In Clusters 7-9, a burst of actin genes marked the formation of a contractile actin ring during cytokinesis, interrupting the two rounds of meiotic nuclear division (**Fig. S7**).
(6) Genotyping of individual cells by SNPs in their RNA reads. This revealed two parallel trajectories (one for each mating type), and pinpointed the timing of cell fusion to the start of Cluster 14 (**Fig. 3a**).
(7) Transcriptional reporter lines. The GEX1 reporter line confirmed the identity of gametes (Cluster 10-13), which are the first cell types expressing this gene. The AAE2 reporter

line, on the other hand, pinpointed the start of auxospore expansion to the intersection between Cluster 14 and Cluster 15. Myb reporter lines supported the position of gametogenesis / gametes.

### Bibliography


1. Roberts, W.R., Ruck, E.C., Downey, K.M., Pinseel, E., and Alverson, A.J. (2023). Resolving Marine–Freshwater Transitions by Diatoms Through a Fog of Gene Tree Discordance. Syst. Biol., syad038. https://doi.org/10.1093/sysbio/syad038.

2. Bilcke, G., Campese, L., Annunziata, R., Amadei Martínez, L., Borgonuovo, C., Rijsdijk, N., Chaerle, P., Van den Berge, K., D'hondt, S., Iudicone, D., et al. (2025). Conserved genetic markers reveal widespread diatom sexual reproduction in the global ocean. Nat. Commun. *16*, 10029.

3. Grujčić, V., Saarenpää, S., Sundh, J., Sennblad, B., Norgren, B., Latz, M., Giacomello, S., Foster, R.A., and Andersson, A.F. (2024). Towards high-throughput parallel imaging and single-cell transcriptomics of microbial eukaryotic plankton. PLOS ONE *19*, e0296672. https://doi.org/10.1371/journal.pone.0296672.

4. Nadal-Ribelles, M., Solé, C., de Nadal, E., and Posas, F. (2024). The rise of single-cell transcriptomics in yeast. Yeast Chichester Engl. *41*, 158–170. https://doi.org/10.1002/yea.3934.

5. Onsbring, H., Tice, A.K., Barton, B.T., Brown, M.W., and Ettema, T.J.G. (2020). An efficient single-cell transcriptomics workflow for microbial eukaryotes benchmarked on Giardia intestinalis cells. BMC Genomics *21*, 448. https://doi.org/10.1186/s12864-020-06858-7.
